# Temporal Coordination of the Transcription Factor Response to H_2_O_2_ stress

**DOI:** 10.1101/2023.03.07.531593

**Authors:** Elizabeth Jose, Woody March-Steinman, Bryce A. Wilson, Lisa Shanks, Andrew L. Paek

**Author notes:** Corresponding author: 1007 E. Lowell Street, PO Box 210106, Tucson, Arizona 85721.

## Abstract

The p53 and FOXO transcription factors (TFs) share many similarities despite their distinct evolutionary origins. Both TFs are activated by a variety of cellular stresses and upregulate genes in similar pathways including cell-cycle arrest and apoptosis. Oxidative stress from excess H_2_O_2_ activates both FOXO1 and p53, yet whether they are activated at the same time is unclear. Here we found that cells respond to high H_2_O_2_ levels in two temporal phases. In the first phase FOXO1 rapidly shuttles to the nucleus while p53 levels remain low. In the second phase FOXO1 exits the nucleus and p53 levels rise. We found that other oxidative stress induced TFs are activated in the first phase with FOXO1 (NF-κB, NFAT1), or the second phase with p53 (NRF2, JUN) but not both following H2O2 stress. The two TF phases result in large differences in gene expression patterns. Finally, we provide evidence that 2-Cys peroxiredoxins control the timing of the TF phases in response to H_2_O_2_.

## Introduction

Hydrogen peroxide (H_2_O_2_) is a reactive oxygen species with a complex role in cellular physiology. H_2_O_2_ is produced as a byproduct of cellular respiration and by over 40 enzymes in humans.^1^ H_2_O_2_ functions as a second messenger, activating receptor tyrosine kinases that promote proliferation, differentiation, and wound healing.^2–6^ Yet at high concentrations, H_2_O_2_ is toxic to cells due to the creation of hydroxyl radicals by the Fenton reaction. Hydroxyl radicals cause DNA damage, lipid peroxidation, and the formation of unfolded/aggregated proteins all of which inhibit cell proliferation and can induce cell death^7,8^. Thus, H_2_O_2_ levels must be tightly regulated and rapidly cleared when concentrations are too high as elevated levels of H_2_O_2_ are thought to be the underlying cause of many human pathologies^9^.

To counter high levels of H_2_O_2_, metazoans activate several transcription factors (TFs) including NRF2, p53, FOXO and other TFs, which act to restore the redox state of the cell and repair damage caused by oxidative stress^10^. Upon activation by H_2_O_2_, these TFs upregulate hundreds of target genes in diverse cytoprotective processes including cell-cycle arrest, NADPH/GSH production, ROS scavenger enzymes, DNA damage repair, autophagy, and protein quality control^11–17^. In addition, both p53 and FOXO can induce cell death by upregulating apoptotic genes^18,19^. Given the diverse molecular challenges caused by oxidative stress, which TFs are activated, and their order of activation is likely tightly regulated and dependent on H_2_O_2_ concentration^20^. Indeed, oxidative stress is often differentiated broadly into either eustress (mild oxidative stress) or distress (toxic oxidative stress) and it is thought that these different levels of stress activate different TFs^1,21^. Yet which TFs are activated at low vs. high oxidative stress and the relative timing of TF activation is not known and is essential for understanding how cells combat oxidative stress and how this might break down in disease.

Evidence from yeast suggests that which TFs are activated, and the order of activation is tightly regulated and determined by the concentration of H_2_O_2_. This is best illustrated by studies of two H_2_O_2_ induced TFs in fission yeast, Pap1 and Atf1. Pap1 is activated rapidly (∼5 minutes) at low levels of H_2_O_2_, while Atf1 remains largely inactive^22,23^. Higher concentrations of H_2_O_2_ activate both Atf1 and Pap1, yet there is a delay (∼30 minutes) in Pap1 activation^24,25^. Why is there immediate activation of Pap1 at low concentrations but a delay at higher concentrations? Pap1 is activated by Tpx1, a 2-Cys peroxiredoxin (PRDX) protein. At low H_2_O_2_ concentrations Tpx1 activates Pap1 through a redox relay mechanism whereby oxidative equivalents stemming from H_2_O_2_ are passed from Tpx1 to Pap1.^26^ Oxidized Tpx1 also acts as a competitive inhibitor to thioredoxin thus blocking reduction of Pap1^27,28^. This leads to an intramolecular disulfide bond in Pap1 which blocks Pap1’s interaction with the nuclear export protein Crm1, causing nuclear accumulation of Pap1 and activation of downstream target genes^29,30^. Yet high concentrations of H_2_O_2_ inactivate Tpx1 due to hyperoxidation of a key cysteine residue in Tpx1 and this prevents Pap1 activation. Hyperoxidation of Tpx1 can be reversed by sulfiredoxin (SRX1 in yeast, SRXN1 in humans), but this takes time, and thus Pap1 activation is delayed until SRX1 repairs hyperoxidized Tpx1^30^.

There is strong evidence that PRDX dependent redox relays also occur in mammals. A knockout model of PRDX1 and PRDX2 in HEK293T cells showed reduced protein disulfide bond formation following oxidative stress, and transient disulfide bond intermediates were recovered between both PRDX1/2 and dozens of other proteins^31^. Further evidence supports a role for PRDX proteins in regulating transcription factors. For example, PRDX2 regulates STAT3 by a redox relay resulting in a disulfide bond in STAT3 causing oligomer formation and attenuation of transcription^32,33^. PRDX1 can form disulfide bonds with FOXO3, which leads to its retention in the cytosol^34,35^. Yet the role of PRDX proteins in regulating the timing of TF activation in response to H_2_O_2_ and how the timing of activation is affected by dose is unclear.

In this study we found that the set of TFs activated by H_2_O_2,_ and their time of activation is dose dependent. We first focused on FOXO1 and p53 as both are activated by H_2_O_2_ and upregulate genes in overlapping pathways including cell-cycle arrest and apoptosis. Using immunofluorescence and time-lapse imaging we found that low levels of H_2_O_2_ cause an immediate increase in p53 levels while FOXO1 remains inactive in the cytoplasm. At higher H_2_O_2_ concentrations there are two temporal phases of activation: in the first phase FOXO1 is shuttled into the nucleus within 1 hour, while p53 levels are kept low. In the second phase, FOXO1 exits the nucleus which is followed by an increase p53 levels. The duration of the first phase, where FOXO1 is active and p53 inactive, increases with H_2_O_2_ dose. Furthermore, we found that other TFs are activated either with FOXO1 (NF-κB, NFAT1) or with p53 (NRF2, JUN) but not both, suggesting coordinated regulation of each group of TFs. The difference in TF activation between the two temporal phases is reflected in large differences in gene expression with increases in ribosome, oxidative phosphorylation, and proteasome genes in phase 1 and NRF2 and p53 target genes involved in NADPH, glutathione, and nucleotide production increasing in phase 2. Finally, we found evidence that the peroxiredoxin/sulfiredoxin system controls which group of transcription factors is activated. The distinct target genes activated in each phase, coupled with the evolutionary conservation of a PRDX control mechanism, suggests that ordering the transcriptional response to H_2_O_2_ is critical for properly restoring redox balance.

## Results

### Mutually exclusive activation of FOXO1 and p53 in response to H_2_O_2_

To determine if FOXO1 and p53 are activated at the same level of H_2_O_2_ stress, we performed a dose response in MCF7 cells and immunostained for FOXO1 and p53 five hours after treatment. FOXO1 is regulated by nuclear/cytoplasmic shuttling, so we measured FOXO1 activation by determining the fraction of nuclear FOXO1 in individual cells^36^. For p53 we measured mean nuclear levels as p53 is predominantly localized to the nucleus and increases due to inhibition of proteasomal degradation.^37^ At low H_2_O_2_ concentrations nuclear FOXO1 levels were unchanged yet p53 levels increased (Figure 1A-C, 20-60 μM). At higher H_2_O_2_ concentrations we observed two distinct populations of cells: one population with increased p53 levels and cytoplasmic FOXO1, and a second population with predominantly active (nuclear) FOXO1 and low p53 levels (Figure 1A-C, 80-100 μM, Figure S1B for activation thresholds). The proportion of cells with nuclear FOXO1 increased with H_2_O_2_ concentration, while cells with high p53 levels decreased. At the highest H_2_O_2_ dose (200 μM), most cells had active FOXO1, while p53 active cells were comparable to untreated controls. Cells with activation of both FOXO1 and p53 were <5% in all doses tested, suggesting that activation of FOXO1 and p53 is mutually exclusive in response to H_2_O_2_.

**Figure 1:**
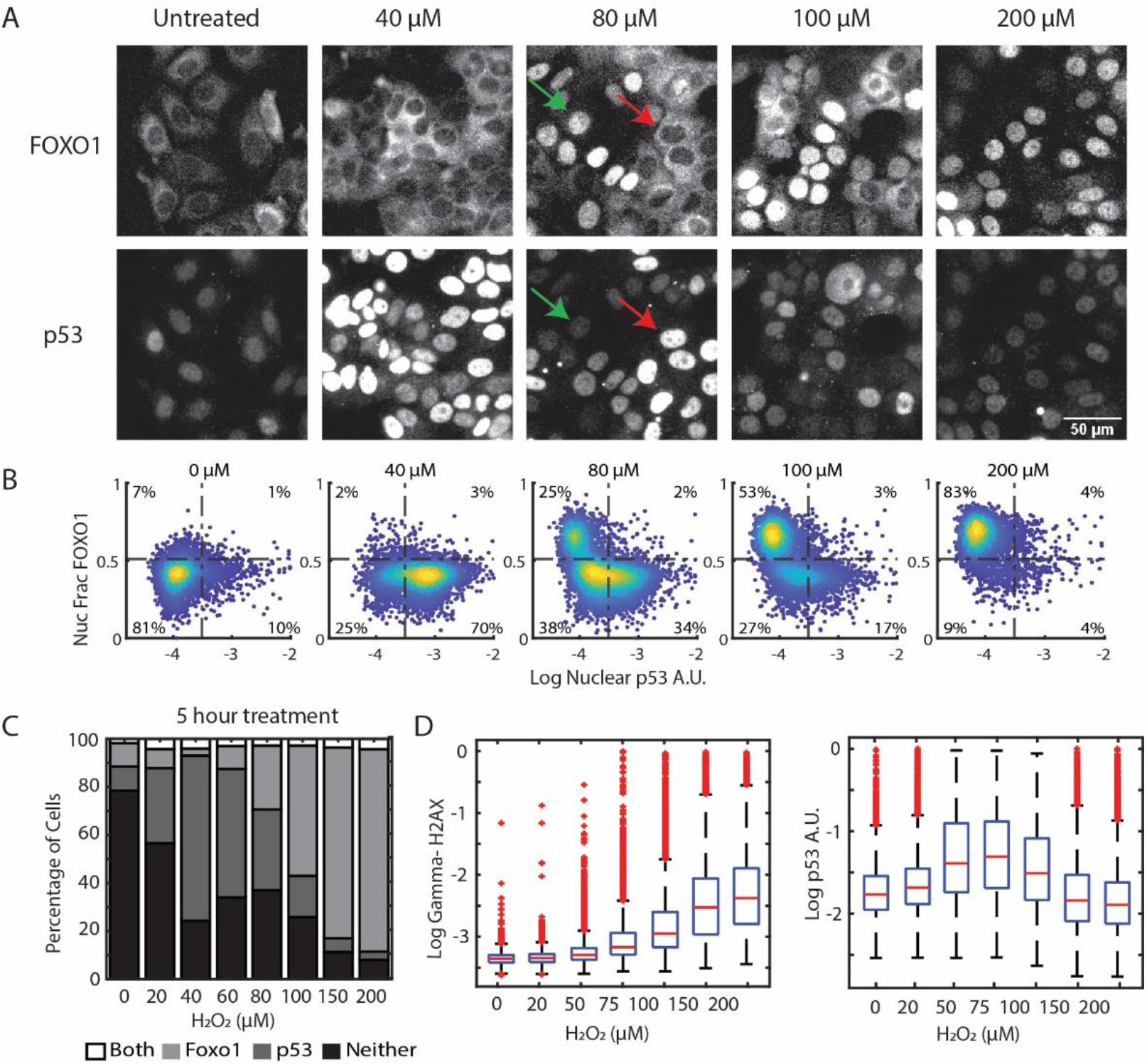
Activation of FOXO1 and p53 in response to H_2_O_2_ is mutually exclusive. (A) Immunofluorescence images of MCF7 cells treated with indicated concentrations of H_2_O_2_ for 5 hours and stained for FOXO1 (top row) and p53 (bottom row). Green arrow indicates a cell with nuclear FOXO1 and low levels of p53. Red arrow shows a cell with cytoplasmic FOXO1 and increased levels of p53 (B) Density colored scatter plots (n ≥ 2000 cells) of the log of nuclear p53 levels (x-axis) and the nuclear fraction of FOXO1 (y-axis). (C) Percentage of cells activating both FOXO1 and p53, only FOXO1, only p53 and neither at the indicated concentrations at 5 hours of H_2_O_2_ treatment. Thresholds are as indicated by dashed lines in B (D) Bar and whisker plots of γH2AX and nuclear p53 levels measured by immunofluorescence after 3 hours of H_2_O_2_ treatment at indicated concentrations. γH2AX and nuclear p53 levels were measured in the same experiment.

The response to H_2_O_2_ is known to depend on the number of cells in an experiment, and indeed we observed the concentration of H_2_O_2_ required to activate FOXO1 increased with the number of cells plated (Figure S1A). Therefore, for all experiments in this study we took care to plate equal numbers of cells in control and treatment groups to properly measure relative activation of each transcription factor.

The lack of p53 activation at higher doses of H_2_O_2_ was unexpected as H_2_O_2_ causes DNA damage, which in principle should activate p53^38^. To ensure that H_2_O_2_ was inducing DNA damage, we repeated the H_2_O_2_ dose-response and measured p53 and γH2AX, a marker for DNA-damage, in single cells using immunofluorescence. As expected, H_2_O_2_ induced a dose-dependent increase in γH2AX, with the highest dose (200 μM) causing the largest increase in DNA damage (Figure 1D). In contrast, p53 levels increased the most at intermediate doses of H_2_O_2_ (50-100 μM) but were comparable to untreated controls at doses >= 150 μM. The amount of DNA damage at higher doses of H_2_O_2_ was comparable to treatment with Neocarzinostatin (NCS), which causes DNA double-strand breaks and upregulates p53, suggesting a control mechanism to actively suppress p53 activation at high doses of H_2_O_2_ (Figure S1C).

Mutually exclusive activation of FOXO1 and p53 in response to H_2_O_2_ is not limited to MCF7 cells as we observed the same pattern in MCF10A, A549 and U2OS cell lines (Figure S1D-F). The oxidative stress inducing agent menadione also induces mutually exclusive activation of FOXO1 and p53 (Figure S1G). However, tert-butyl hydroperoxide activated p53 but not FOXO1 suggesting a different mode of activation (Figure S1H). Menadione induces formation of superoxide radicals which then dismutate to H_2_O_2_, it is possible that mutually exclusive activation of p53 and FOXO1 is specific to H_2_O_2_. Together these data suggest that mutually exclusive activation of p53 and FOXO1 is not cell-type specific but does not occur under all forms of oxidative stress.

### FOXO1 activation precedes p53 activation at high concentration of H_2_O_2_

The surprising lack of p53 activation at high H_2_O_2_ concentrations might be due to a temporal delay in activation as our immunofluorescence experiments were performed five hours after H_2_O_2_ treatment. To test this hypothesis, we tagged FOXO1 and p53 genes with fluorescent reporters to measure both TFs in single cells in real time. We used a previously developed MCF7 cell line where CRISPR/Cas9 was used to tag the endogenous locus of FOXO1 at the C-terminus with the mVenus fluorescent protein. This cell line also contains a H2B-CFP tag for tracking nuclei^39^. For p53, we added an exogenously expressed p53-mCherry reporter used and validated in previous studies^40^. We then treated cells containing both reporters with four different concentrations of H_2_O_2_ and measured TF levels in single cells every 20 minutes for 24 hours. At the lowest H_2_O_2_ dose (50 μM), FOXO1 remained largely inactive (in the cytoplasm), while p53 levels oscillated, as shown previously in response to DNA double strand breaks and H_2_O_2_ ^40,41^(Figure 2A, 2B, Supp Movie 1).

**Figure 2:**
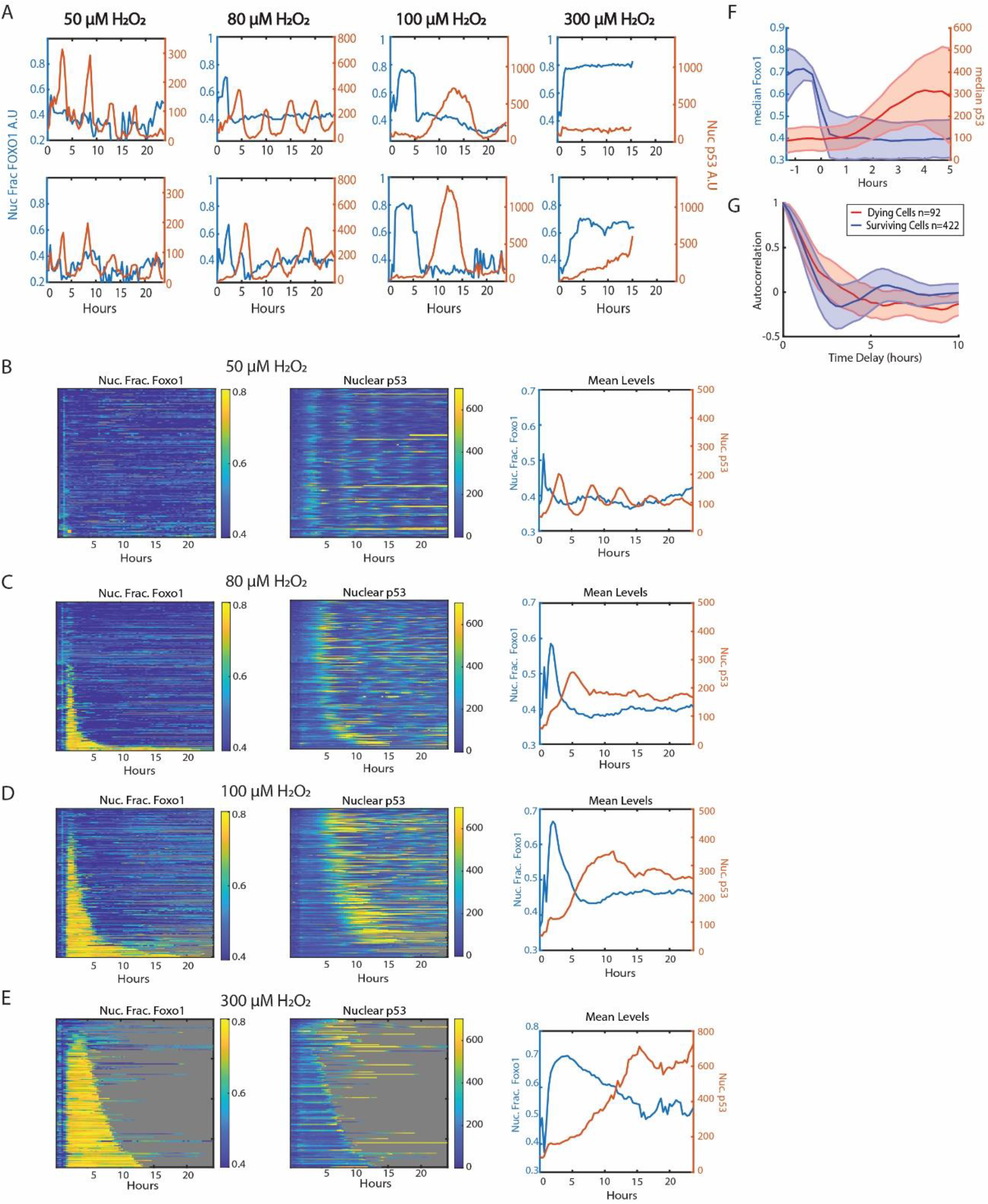
Live cell imaging of FOXO1 and p53 over time shows sequential activation of FOXO1 followed by p53. (A) Sample single cell traces of nuclear fraction of FOXO1 (blue, left y-axis) and nuclear levels of p53 (red, right y-axis) of cells treated with 50μM, 80μM, 100μM and 300μM of H_2_O_2_ for 24 hours. (B-E) Heat maps of single-cell traces of nuclear fraction of FOXO1-mVenus (left), nuclear p53-mCherry (middle), and mean of both (right) for 24 hours following H_2_O_2_. Each row of the heat maps is a single cell over time. Both FOXO1 and p53 heat maps are sorted by the duration that FOXO1-mVenus remained in the nucleus. Gray indicates cell death. (B) 50μM H_2_O_2_ (n=196 cells, 1% cell death). (C) 80μM H_2_O_2_ (n=250 cells, 11% cell death). (D) 100μM H_2_O_2_ (n=300 cells, 34% cell death). (E) 300μM H_2_O_2_ (n=206 cells, 98% cell death). (F) Median levels of FOXO1 and p53 of all cells in 80μM and 100μM treatments with traces aligned to when FOXO1 exited the nucleus. (G) Autocorrelation of p53 trajectories of dying (red, n=92) vs surviving (blue, n=422) cells, treated with 80μM and 100μM of H_2_O_2_ from Figures 2C and 2D.

At higher H_2_O_2_ concentrations, FOXO1 accumulated in the nucleus within 1 hour of treatment in a subset of cells (Figure 2A,C-E, Supp Movies 2,3,4). While FOXO1 was in the nucleus, p53 levels remained low, and only increased after FOXO1 exited the nucleus. The fraction of cells with nuclear FOXO1 and the time in which FOXO1 remained in the nucleus increased with H_2_O_2_ dose. To visualize data from multiple cells, we created single-cell heat maps of nuclear FOXO1 and p53, where cells were sorted by the time in which FOXO1 remained in the nucleus (Figure 2B-E). Aligning trajectories to the time that FOXO1 exited the nucleus revealed that p53 began accumulating ∼1 hour after FOXO1 exited the nucleus (Figure 2F). Together these data revealed that there are two temporal phases to the FOXO1/p53 response to high concentrations of H_2_O_2_. In the initial phase (phase 1), FOXO1 enters the nucleus and p53 levels remain low. In the second phase (phase 2), FOXO1 exits the nucleus and p53 begins to accumulate within 1 hour.

The dynamics of p53 accumulation also differs in response to higher concentrations of H_2_O_2_. In some cells p53 levels oscillate similar to the 50 μM dose, yet often with a higher initial spike in p53 levels (Figure 2A). While other cells show large bursts of p53 levels similar to the response to UV irradiation^42^. The proportion of cells with oscillating p53 levels decreased with dose as shown by autocorrelation analysis, and recently observed in retinal pigment epithelial cells ^40^(Figure S2A-D). These differences correlated with cell survival as the p53 levels in cells that died reached higher levels than those that survived (Figure S2G and S2H). In addition, autocorrelation analysis revealed oscillations in p53 activation in surviving cells but not in dying cells (Figure 2G). Dying cells also showed an increase in the duration of FOXO1 activation as compared to surviving cells, in agreement with our previous study ^39^(Figure S2E,F). Together these data show that higher concentrations of H_2_O_2_ cause prolonged activation of FOXO1, a delayed yet stronger p53 response, and an increase in cell death.

### Additional H_2_O_2_ induced transcription factors are activated with either FOXO1 or p53

The activation of FOXO1 and p53 in two distinct temporal phases prompted us to ask whether other transcription factors are activated with FOXO1 in phase 1, or p53 in phase 2 in response to H_2_O_2_. To identify potential transcription factors, we treated MCF7 cells with PBS as a control and two different concentrations of H_2_O_2_ (50μM and 75μM), isolated individual nuclei, and performed single-cell Assay for Transposase-Accessible Chromatin using sequencing (ATAC-seq) and gene expression using the 10X genomics single-cell Multiome kit. Unsupervised clustering of the ATAC-seq data identified six separate clusters (Figure 3A and 3B). Clusters four and five represented PBS control nuclei, while the other clusters were observed predominantly in H_2_O_2_ treated nuclei. Clusters two and three were enriched with FOXO motifs but not p53 motifs and we refer to these clusters together as the FOXO cluster (Figure S3A). In contrast, nuclei in cluster six were enriched for p53 motifs, but not FOXO motifs (Figure S3B). Thus, we can capture cells in phase 1 (FOXO1 active, p53 inactive) and phase 2 (p53 active, FOXO1 inactive) by ATAC-seq.

**Figure 3:**
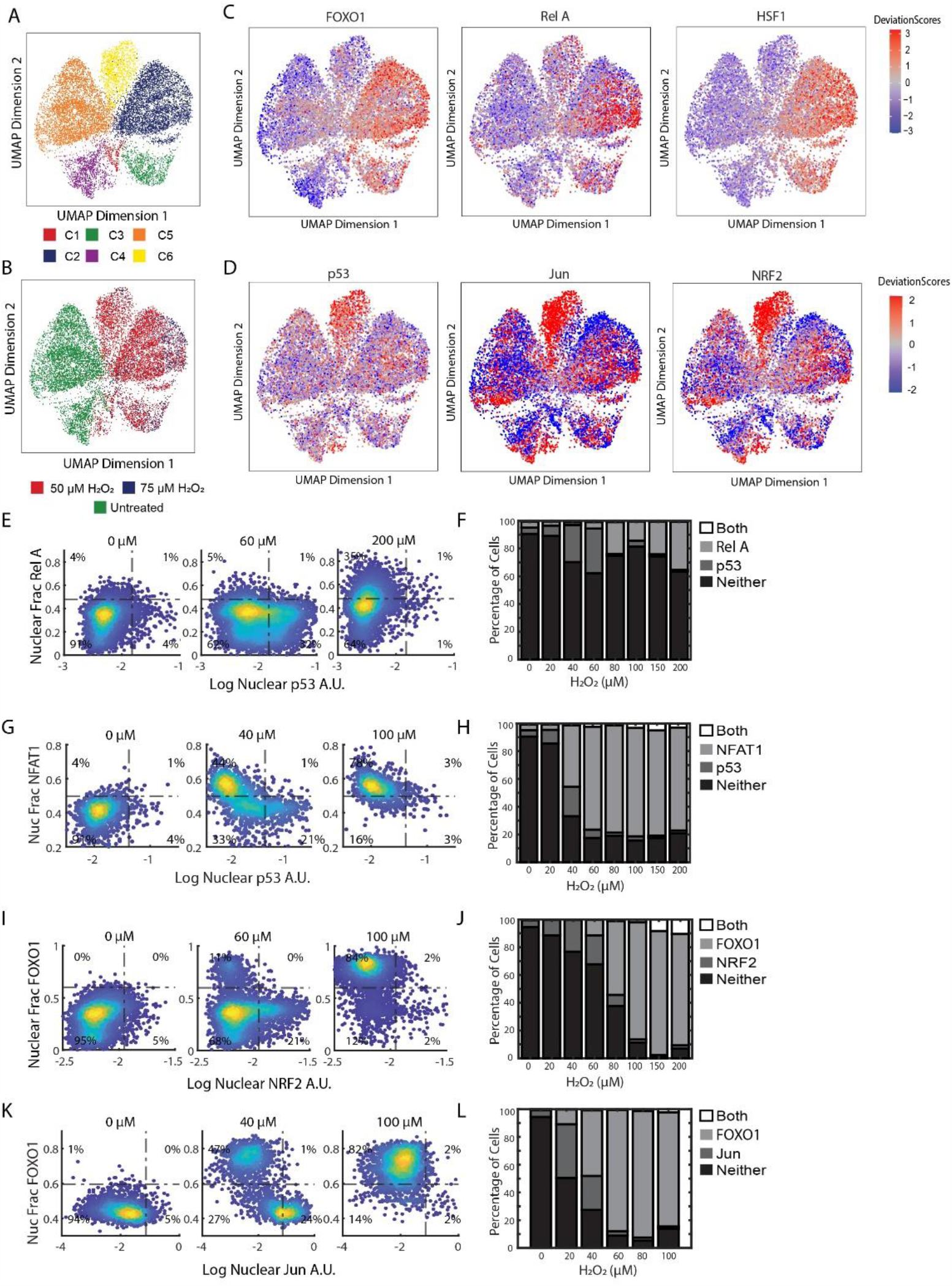
Single-cell ATAC-seq reveals transcription factors activated with FOXO1 or p53 in response to H_2_O_2_. (A) Uniform manifold approximation and projection (UMAP) plot of cells treated with PBS, 50μM and 75μM of H_2_O_2_ after unsupervised clustering (n ≥ 10,000). Colors for cells based on the six clusters obtained. (B) UMAP of the same cells in A but colored based on sample. (C) UMAPs of cells from A colored by deviation scores for FOXO1 (left), RelA (middle) and HSF1 motifs (right) (D) UMAPs of cells from A showing deviation scores for p53 (left), Jun(middle) and NRF2 motifs (right) (E) Density colored scatter plots of log nuclear p53 (x-axis) and nuclear fraction of RelA (y-axis) at indicated levels of H_2_O_2_ treatment for 5 hours (F) Percentage of cells activating both RelA and p53 (Both), RelA only, p53 only or neither for all concentrations of H_2_O_2_ (G) Density colored scatter plots of log nuclear p53 (x-axis) and nuclear fraction of NFAT1 (y-axis) at indicated levels of H_2_O_2_ treatment for 5 hours (H) Percentage of cells activating both NFAT1 and p53 (Both), NFAT1 only, p53 only or neither for all concentrations of H_2_O_2_. (I) Density colored scatter plots of log nuclear NRF2 (x-axis) and nuclear fraction of FOXO1(y-axis) at indicated levels of H_2_O_2_ treatment for 5 hours (J) Percentage of cells activating both FOXO1 and NRF2 (Both), FOXO1 only, NRF2 only or neither for all concentrations of H_2_O_2_. (K) Density colored scatter plots of log nuclear JUN (x-axis) and nuclear fraction of FOXO1 (y-axis) at indicated levels of H_2_O_2_ treatment for 5 hours (L) Percentage of cells activating both FOXO1 and JUN (Both), FOXO1 only, JUN only or neither for all concentrations of H_2_O_2_.

We then focused on other transcription factor motifs that were enriched with either the FOXO or p53 cluster but not both (Figure S3A and S3B). Within the FOXO cluster, we observed an enrichment in HSF, GRHL1, NF-KB, ZKSCAN1 and NFAT TF motifs (Figure 3C and S3A). To determine if these TFs showed similar activation kinetics as FOXO1, as suggested by the ATAC data, we measured the NF-kB isoform RELA and NFAT isoform NFAT1 together with p53 in single cells by immunofluorescence 5 hours after H_2_O_2_ treatment (Figures 3E-H). Similar to FOXO1, we observed mutually exclusive activation of RELA and NFAT1 with p53, with both TFs showing maximum activation at the highest concentration of H_2_O_2_ (Figure 3F and 3H). RELA was only activated at high concentrations of H_2_O_2_ and only in a subset of cells consistent with the ATAC data (Figure 3C). We also measured nuclear HSF1 levels, but did not observe a change in concentration in response to H_2_O_2_ (Figure S3D). Thus the enrichment in HSF motifs might be due to upregulation of HSF2/HSF4, or regulation of these factors independent of an increase in nuclear levels as has been described elsewhere^43^. Cells in the FOXO clusters (clusters 2 & 3) did have an increase in the expression of the canonical HSF target genes HSPA1A (log2 fold-change .98, P <10^-5) and HSPA1B log2 fold-change 1.1, P <10^-6), consistent with a role of heat shock protein activation during the FOXO phase.

The p53 cluster was enriched for motifs in the AP-1 family of TFs (JUN, FOS, and ATF subfamilies), as well as NRF2 (*NFE2L2* gene) (Figure 3D and Figure S3B). The AP-1 family have similar binding motifs which might give false positives for activation^44^. We picked two AP-1 transcription factors (JUN and FOS) and NRF2 to validate the ATAC data. JUN and NRF2 showed similar patterns to p53 activation (Figures 3I and 3K), increasing at low concentrations of H_2_O_2_ and little to no activation at higher concentrations (Figures 3J & 3L). FOS on the other hand was activated by H_2_O_2_ in cells with both nuclear FOXO1 and cytoplasmic FOXO1 (Figure S3A). Together these data suggest that other H_2_O_2_ activated transcription factors are upregulated either with FOXO1 (RELA, NFAT1) or p53 (JUN, NRF2), but not both, following H_2_O_2_ treatment.

### The role of the Peroxiredoxin/Sulfiredoxin system in controlling the switch between p53 and FOXO1 activation

Next, we investigated the mechanism underlying the switch from active p53 and inactive FOXO1 at low H_2_O_2_ concentrations, to active FOXO1 and inactive p53 at high H_2_O_2_ concentrations. Since other transcription factors are activated with either FOXO1 or p53, with distinct mechanisms of control, it is likely that the mechanism lies upstream of these factors. A redox relay stemming from a 2-Cys PRDX protein is a plausible mechanism as redox relays can affect multiple proteins and are switched off by hyperoxidation when H_2_O_2_ levels cross a particular threshold, as described below.

2-Cys PRDX proteins function as dimers and harbor two key cysteine residues: a peroxidatic cysteine (Cp), whose thiol group is oxidized to sulfenic acid (SOH) by H_2_O_2_, and a resolving cysteine (Cr), which reacts with the sulfenic acid form of Cp in trans, forming a disulfide bond between the two monomers ^45^(Figure 4A). This disulfide bond can be resolved by the thioredoxin system, or it can participate in a redox relay, where it transfers oxidative equivalents to downstream proteins, altering their function^46^. However, at high levels of H_2_O_2_, the Cp of PRDX is hyperoxidized from sulfenic acid to sulfinic acid (SO_2_H), rendering it incapable of forming a disulfide bond with Cr or participating in redox relays (Figure 4A). We hypothesized that the switch from p53 activation at low H_2_O_2_ levels to FOXO1 activation at high H_2_O_2_ levels is due to hyperoxidation of a PRDX protein. Switching off FOXO1 and the subsequent activation of p53 would only occur after SRXN1 repairs the hyperoxidized PRDX protein, similar to Pap1 activation in yeast (Figure 4A).

**Figure 4:**
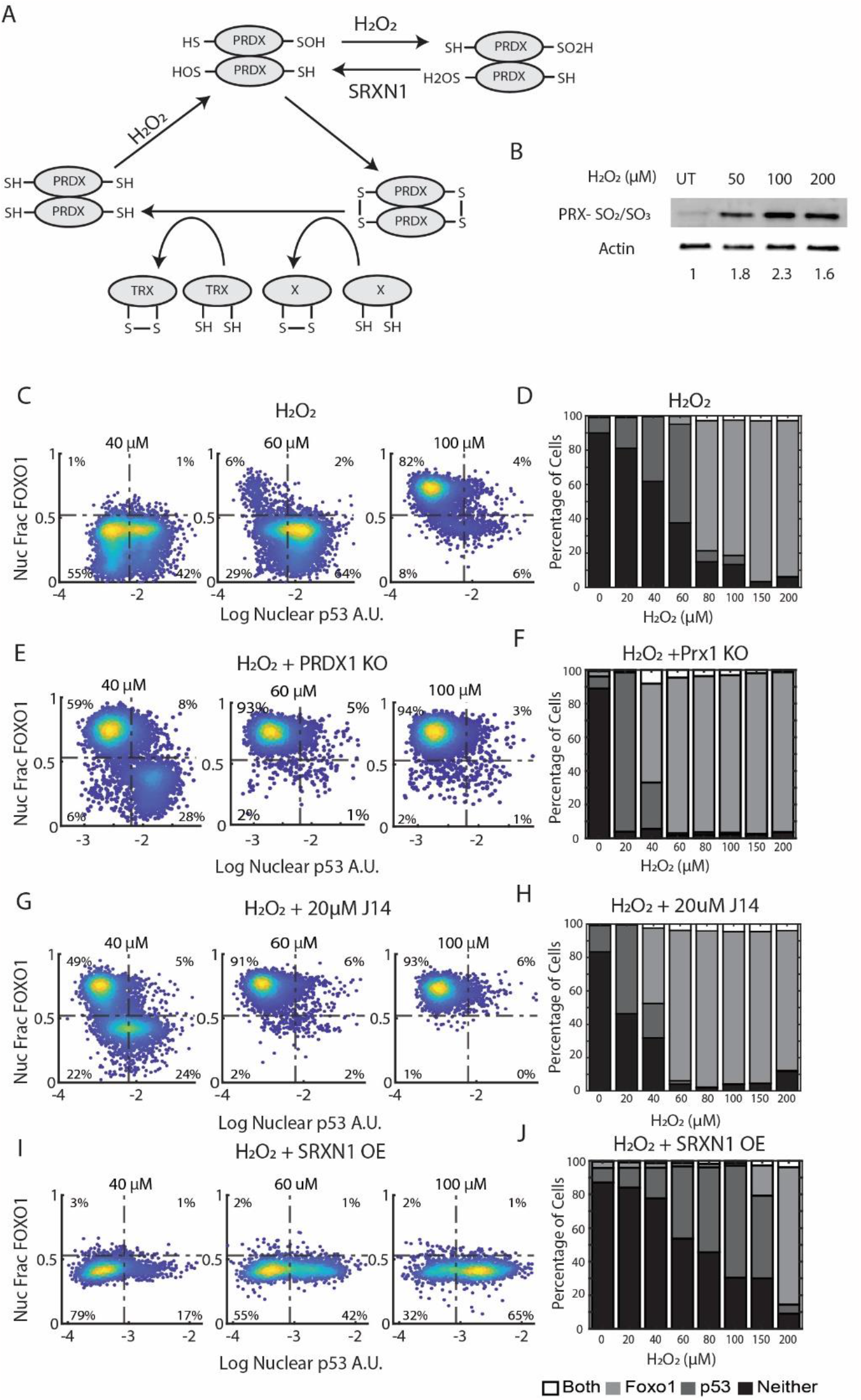
The Peroxiredoxin/ Sulfiredoxin system sets the threshold for p53 and FOXO1 activation. (A) Schematic of the redox cycle of PRDXs. H_2_O_2_ oxidizes the peroxidatic cysteine to sulfenic acid (SOH), which can form a disulfide bond in trans with the resolving cysteine. At high H_2_O_2_ concentrations, the peroxidatic cysteine is further oxidized (hyperoxidized, SO2H). Hyperoxidized PRDX can be repaired by SRXN1 to sulfenic acid. Oxidized peroxiredoxins can be reduced by the Thioredoxin system or can transfer oxidative equivalents to other proteins like i.e. protein X in the diagram. (A) Western Blot stained for hyperoxidized (SO_2_/SO_3_) PRDX1/2 and Actin of MCF7 cells treated with indicated concentrations of H_2_O_2_ for 3 hours. Density colored scatter plots (C, E, G & I) and percentage of cells activating both FOXO1 and p53, only FOXO1, only p53 or neither (D, F, H & J). (C & D) Cells treated with H_2_O_2_ used as a control at the indicated concentrations of H_2_O_2_ for 5 hours. (E & F) PRDX1 knockout cells treated with H_2_O_2_ at indicated concentrations for 5 hours. (G & H) Cells treated with 20μM of J14, an inhibitor of Sulfiredoxin, along with H_2_O_2_ at the indicated concentrations for 5 hours. (I & J) Cells overexpressing Sulfiredoxin (SRXN1-OE) treated with mentioned concentrations of H_2_O_2_ for 5 hours.

To test this hypothesis, we first measured hyperoxidation of PRDX1/2 following H_2_O_2_ treatment by western blot. We observed dose-dependent hyperoxidation at concentrations in which cells activated FOXO1 in previous experiments (Figure 4B). We then tested the effect of Conoidon A, an inhibitor of PRDX1 and PRDX2^47,48^. Treatment of cells with 5 μM Conoidon A resulted in the activation of FOXO1 and prevented p53 accumulation at low doses of H_2_O_2_ (Figure S4A), consistent with a role for hyperoxidation and inactivation of PRDX1 or PRDX2 as key events in activating FOXO1 and repressing p53 activation. Furthermore, 10 μM Conoidon A resulted in FOXO1 activation but not p53 activation in the absence of H_2_O_2_ (Figure S4A). Taken together, these data suggest that inactivation of PRDX1 and/or PRDX2 results in FOXO1 activation and suppression of p53 activation in response to H_2_O_2_.

We then tested the specific role of PRDX1, as it forms disulfide bonds with FOXO proteins, and knockdown of PRDX1 increased nuclear FOXO3 in response to H_2_O_2_ in a previous study^34,35^. We created a PRDX1 knockout line using CRISPR/Cas9 (Figure S4B). In the absence of PRDX1, there is a substantial increase in FOXO1 active cells and a subsequent decrease in p53 active cells at H_2_O_2_ concentrations ≥ 40μM and above (Figures 4E and 4F). There was an increase in p53 activation at 20μM H_2_O_2_ in PRDX1 knockout cells compared to controls. This is likely due to an increase in DNA damage, as PRDX1 is required for proper repair of DNA DSBs^49^. Consistent with this, DNA damage, as measured by γH2AX, was significantly increased in PRDX1 knockout cells following H_2_O_2_ treatment (Supp Figure 4C). The decreased H_2_O_2_ threshold for FOXO1 activation in PRDX1 knockout cells supports a role for PRDX1 inactivation as a key step in activation of FOXO1 and suppression of p53 activation.

To test the role of PRDX hyperoxidation in activating FOXO1 and inhibiting p53 we used J14, a small molecule inhibitor of SRXN1. At concentrations of H_2_O_2_ 40 μM and above, J14 led to a decrease in p53 active cells, and a corresponding increase in FOXO1 activation similar to PRDX1 knockout cells (Figures 4G and 4H). To confirm that the effect of J14 was due to inhibition of SRXN1 and not due to off-target effects, we knocked down SRXN1 using shRNA (Figure S4F). SRXN1 knockdown showed a similar shift from p53 activation to FOXO1 activation (Figure S4D). In addition, live-cell microscopy revealed that J14 prolonged FOXO1 activation and delayed p53 activation, supporting the role of SRXN1 in shutting of FOXO1 and activation of p53 (Supp Figure 4E). Prolonged FOXO1 activation by J14 was accompanied by increased cell death.

If hyperoxidation of PRDX1 is required for FOXO1 activation and p53 repression, SRXN1 overexpression should increase the concentration of H_2_O_2_ required for FOXO1 activation. To test this, we established a cell line (SRXN1-OE) in which SRXN1 is expressed from the PGK promoter and verified that these cells have reduced PRDX1/2 hyperoxidation by western blot (Figure S4H). SRXN1-OE cells showed a dramatic decrease in FOXO1 activity at high concentrations of H_2_O_2_ (80-150 μM) and a corresponding increase in p53 activation (compare Figures 4I and 4J to Figures 4C and 4D). The effect of SRXN1 overexpression was reversed by treatment with the SRXN1 inhibitor, J14 (Figure S4I). Together, these data suggest that oxidation of PRDX1 by H_2_O_2_ activates the p53 pathway at low H_2_O_2_ concentrations. At higher concentrations, PRDX1 is hyperoxidized, which inactivates PRDX1 dependent redox relays, shutting off p53 while activating FOXO1. Repair of PRDX1 hyperoxidation by SRXN1, which occurs after a delay, shuts off FOXO1 and restores p53 activity.

### Evidence that p53 is activated by a separate hyperoxidation event that occurs at low H_2_O_2_ concentrations

As noted above, SRXN1-OE cells activated p53 and inhibited FOXO1, at much higher levels of H_2_O_2_ (80-150 μM) than control cells. However, at low levels of H_2_O_2_ (20-60 μM), SRXN1-OE showed *fewer* p53 active cells than control cells (Figure 4Is and 4J). This suggests that a separate hyperoxidation event, which occurs at low H_2_O_2_ levels, is likely required for activation of p53 in some cells. In support of this, inhibition of SRXN1 with J14 caused an increase in p53 active cells at 20 μM H_2_O_2_ (Figure 4H). We propose that there are two mechanisms for activating p53, hyperoxidation and inactivation of a protein that occurs at low concentrations of H_2_O_2_, and activation through H_2_O_2_ induced DNA damage.

### The two temporal phases of transcription factor activation cause distinct transcriptional changes

We next asked what gene expression changes occur in the two transcription factor phases in response to H_2_O_2_ using RNA-seq. To identify genes activated in each phase, we used three different treatment groups, MCF7 cells, MCF7 cells treated with the SRXN1 inhibitor J14, and MCF7 cells overexpressing SRXN1. We performed RNA-seq on each group with and without 50μM H_2_O_2_ as this concentration led to a mix of FOXO1 and p53 active cells with H_2_O_2_ alone, mostly FOXO1 active cells in the J14 treatment group (phase 1), and mostly p53 active cells in the SRXN1 overexpression group (phase 2). Log2 fold changes (L2FC) and p-values were calculated by comparing H_2_O_2_ treated samples to their relevant PBS controls.

A heatmap of differentially expressed genes revealed broad differences in gene expression between J14 and SRXN1-OE samples (Figure 5A). Gene Set Enrichment Analysis (GSEA) using a list of known p53 and NRF2 target genes verified that these genes were enriched in SRXN1-OE cells when compared to J14 treated cells (Figures S5A and S5B, adjusted P < 1^*^10^−9^). Upregulated genes in the p53 and NRF2 pathway included genes in DNA repair (XPC, POLH, DDB2, ASCC3), pentose phosphate pathway/nucleotide biosynthesis (TIGAR, RRM2B, MTHFD2), NADPH production (ALDH3A1, ME1) and glutathione synthesis (GCLM,GCLC,SLC7A11). Similarly, a volcano plot comparing gene expression in the SRXN1-OE cells to J14 treated cells revealed that SRXN1-OE cells show significant increases in p53 (CDKN1A, MDM2, SESN2) and NRF2 (SLC7A11, TXNRD1, HMOX1) target genes (Figure 5B). While the J14 treated samples show significant increases in heat shock protein genes (HSPA1A, HSPA1B).

**Figure 5:**
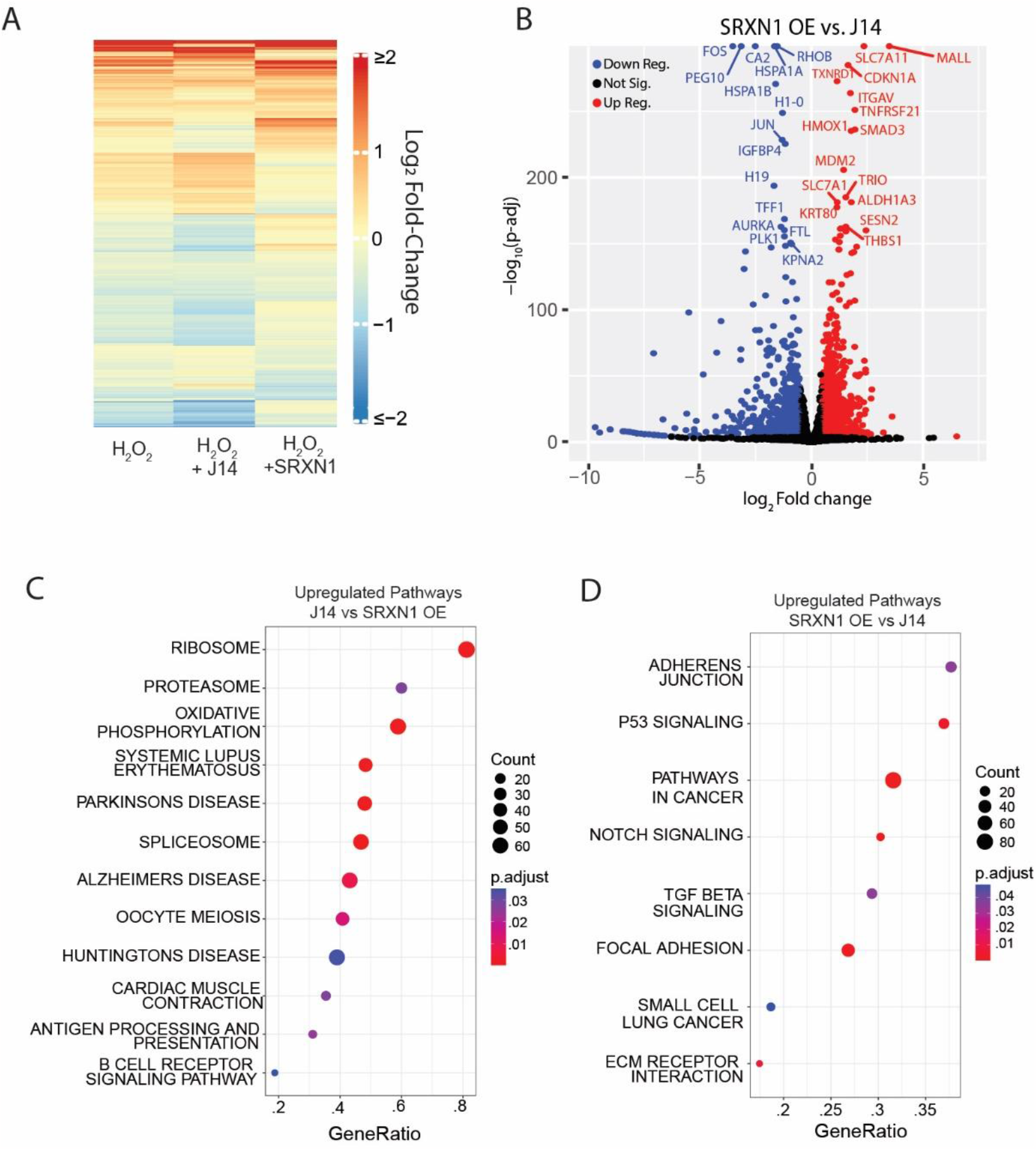
RNA-seq analysis of the two TF phases. (A) Significantly up and downregulated genes (P<.0001, |Log_2_Fold Change| > .2) in H_2_O_2_ treated cells vs. PBS controls, H_2_O_2_ + J14 treated cells vs J14 treated controls and H_2_O_2_ treated SRXN1-OE cells vs. SRXN1-OE PBS controls. H_2_O_2_ concentration is 50μM, J14 20μM. (B) Volcano plot showing log_2_ fold-change and -log_10_ adjusted p-value of SRXN1-OE cells treated with H_2_O_2_ vs J14 treated with H_2_O_2_. (C) GSEA of KEGG pathways upregulated in cells treated with J14 + H_2_O_2_ as compared to SRXN1-OE + H_2_O_2_. (D) GSEA of KEGG pathways upregulated in cells treated with SRXN1-OE + H_2_O_2_ as compared to J14 + H_2_O_2_.

We next focused on the differences between the H_2_O_2_ response in SRXN1-OE and J14 treated cells. GSEA using the hallmark (H), curated gene sets (C2), and regulatory gene sets (C3) from Molecular Signatures Database revealed that J14 treated cells, show significant upregulation of ribosome, proteasome, oxidative phosphorylation, and spliceosome genes when compared to SRXN1-OE cells (Figures 5C and D). Together these data suggest that in phase 1, cells upregulate critical components for protein production (ribosomes, spliceosome), protein quality control (proteasome, heat shock proteins) and oxidative phosphorylation. In contrast during phase 2, cells upregulate genes in DNA repair, the pentose phosphate pathway, nucleotide biosynthesis, NADPH production and glutathione biosynthesis.

## Discussion

Excess levels of H_2_O_2_ activate a diverse set of TFs, including p53, NRF2, JUN, FOXO, NF-KB, and NFAT that act to restore the cellular redox environment and repair cell damage induced by H_2_O_2_. Here we found that which TFs are activated, and their timing of activation are dose dependent. At low levels of H_2_O_2_ stress, cells activate p53, NRF2 and JUN while FOXO, NF-KB and NFAT remain inactive. At high levels of H_2_O_2_ there are two distinct temporal phases of TF activation (Figure 6). In the first TF phase, FOXO, NF-KB and NFAT are activated by shuttling to the nucleus, while p53, NRF2 and JUN remain inactive. In the second phase, FOXO, NF-KB and NFAT switch off and p53, NRF2 and JUN switch on.

**Figure 6:**
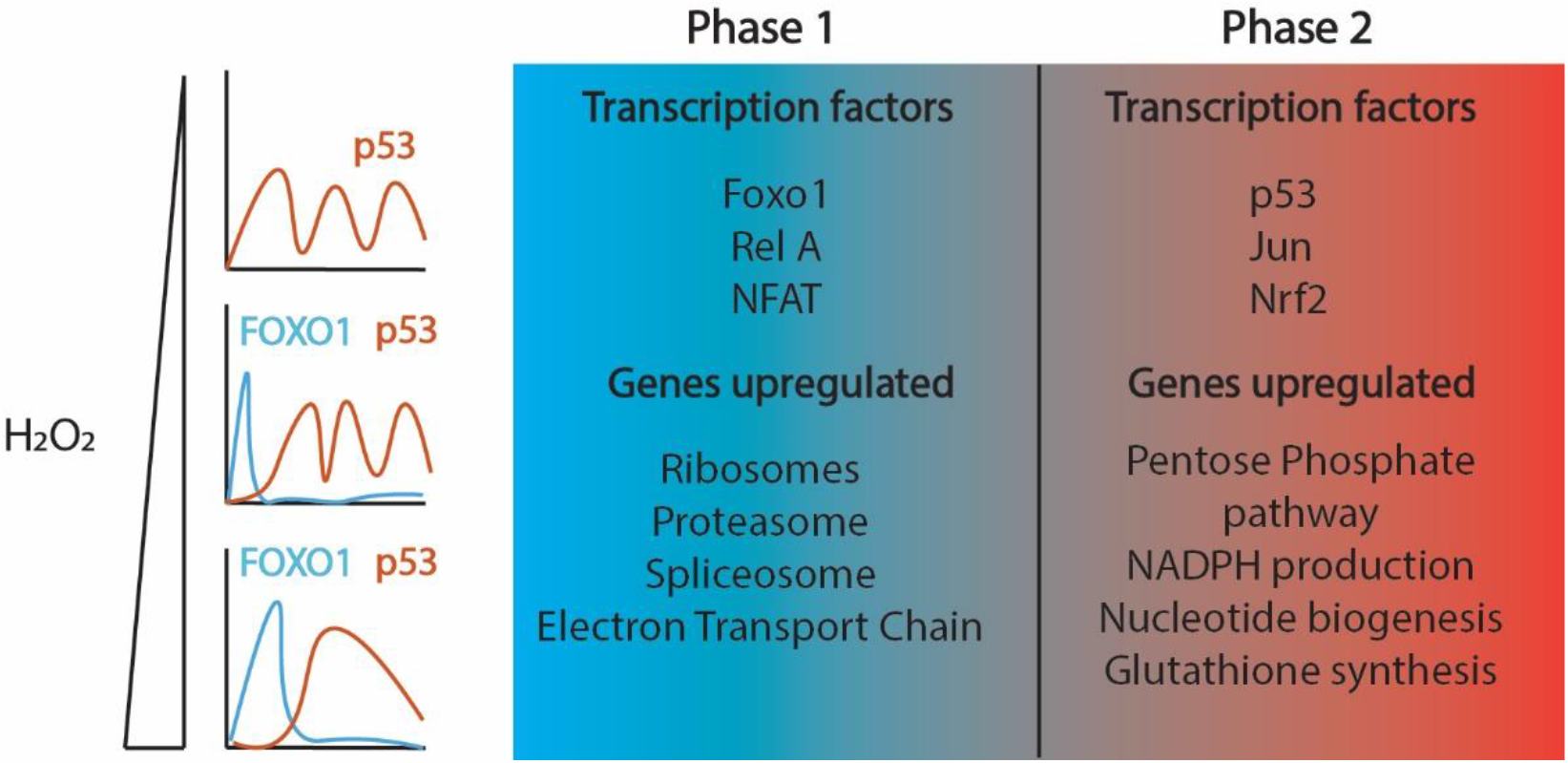
Temporal activation of transcription factors by oxidative stress. Model showing early Foxo1 activation (Phase 1) followed by p53 activation (Phase 2) in response to H_2_O_2_. Lower levels of H_2_O_2_ induce p53 oscillations whereas higher levels of H_2_O_2_ cause p53 to become more sustained in cells. The duration of FOXO1 increases as the dose of H_2_O_2_ is increased. Different transcription factors are activated in each phase. Some are activated earlier along with FOXO1 (Phase 1), indicated in blue, while others are activated later with p53 (Phase 2), indicated in red. The two transcription factor phases upregulate very distinct transcriptional programs.

There are several possible reasons for ordering the TF response at high concentrations of H_2_O_2._ The two different TF phases might set the order of repair of oxidative damage. During the first TF phase there is an increase in expression of ribosome, proteasome, and oxidative phosphorylation genes (components of the electron transport chain). Thus, the initial TF phase might be required for degrading or preventing the formation of protein aggregates, while replacing damaged ribosomes and proteins in the electron transport chain (ETC) for protein and energy production. Indeed, hyperoxidized PRDX proteins assemble into high molecular weight ring structures that function as chaperones with holdase activity^50–52^. In addition, ribosomal RNA and proteins are damaged by oxidative stress and restoring ribosomes might be a high priority for cells immediately following oxidative stress to replace damaged proteins^53^. In a similar fashion, restoring damaged lipids, proteins and DNA/RNA is energy intensive and upregulating ETC genes might be required to ensure adequate energy production. Damage to the ETC can lead to ETC uncoupling, which in turn leads to an increase in H_2_O ^54^ Thus, directly adding H_2_O_2_ to cells might recreate the signal that the ETC is damaged, without damaging components of the ETC.

During the second TF phase there is increased expression of genes involved in NADPH and glutathione synthesis. It is not clear why expression of these genes is delayed at high H_2_O_2_ concentrations. It is possible that the delay in upregulation is required to maintain cysteine oxidation of signaling proteins to maintain activation of TFs in the initial phase. This could ensure adequate induction of genes in the first TF phase, either to ensure proper repair, or alternatively for ensuring cell death if the initial TF phase is prolonged. Indeed, prolonged FOXO activation following H_2_O_2_ treatment is associated with increased cell death^39^. There is also increased expression of genes in the pentose phosphate pathway and nucleotide biosynthesis in the second transcription factor phase. Upregulation of these genes are likely important for completing DNA replication and DNA repair.

We found evidence that the PRDX/SRXN1 system determines which group of TFs are activated, and when following H_2_O_2_ stress. Overexpression of SRXN1, which reduces hyperoxidation of PRDX proteins, increases both the concentration of H_2_O_2_ required to activate p53, and the concentration necessary for switching from p53 to FOXO1 activation. This is consistent with two hyperoxidation events involved in activating p53/FOXO1. One hyperoxidation event happens at low H_2_O_2,_ and activates p53, although some p53 activation is likely due to DNA damage independent of a hyperoxidation event. The second hyperoxidation event, which occurs at high H_2_O_2_ levels leads to FOXO1 activation. We favor hyperoxidation of PRDX1 as the likely event that leads to FOXO1 activation, as PRDX1 knockouts activate FOXO1 at much lower concentrations of H_2_O_2_. *In vitro* measurements of the oxidation rate of different PRDX proteins to sulfenic acid (SOH), the rate of disulfide bond formation, and the rate of oxidation to sulfinic acid (SO_2_H), have revealed substantial differences across the family of PRDX proteins^55–57^. This has prompted others to propose that the different PRDX proteins work to detect and respond to different concentrations of H_2_O_2_ and our data is consistent with this model. However other proteins outside the PRDX family are hyperoxidized by H_2_O_2,_ and are repaired by SRXN1, and we cannot rule out that these PRDX independent hyperoxidation events are involved in activation of p53 of FOXO1^58^. These PRDX independent hyperoxidation events could also be required for activation of p53 or FOXO1.

There is abundant evidence for PRDX dependent redox relays controlling TF activation. In yeast, hyperoxidation of Tsa1 has been shown to delay activation of Pap1. Redox relays can also occur through glutathione peroxidases. In budding yeast, Yap1 activation involves a redox relay from oxidized Gpx3^59^. In addition, budding yeast strains lacking all five peroxiredoxin genes and all 3 glutathione peroxidases, were incapable of altering their transcriptional response to H_2_O_2_, suggesting these proteins are critical for the TF response to H_2_O_2_.

The direct targets of the redox relays that regulate the TFs in this study are not known although many of the TFs in this study, or their upstream regulators, are known to harbor cysteines that regulate their activity. For example, KEAP1 regulates NRF2 by sequestering it in the cytoplasm and targeting it for ubiquitination by the BTB-CUL3-RBX1 E3 ubiquitin ligase complex^60^. H_2_O_2_ treatment leads to a disulfide bond between two cysteines in KEAP1 that prevent NRF2 degradation^61^. Activation of p53 in response to H_2_O_2_ has been linked to ATM, p38 and JNK kinase activity and all three kinases are known to be activated by disulfide bonds either on the protein itself (ATM) or in upstream proteins which regulate kinase activity (p38, JNK)^18,62–64^. Similarly FOXO3 has been shown to form disulfide bonds with PRDX1 which sequester FOXO3 in the cytoplasm^34,35^. In addition, AKT, which phosphorylates FOXO proteins causing cytoplasmic sequestration, is inhibited by disulfide bond formation between two cysteines in the protein, which leads to its inactivation and nuclear accumulation of FOXO^65,66^. In future studies, it will be interesting to elucidate the exact steps of these redox relays.

## Materials and Methods

### Cell Culture

MCF7 (gift from Galit Lahav, Harvard Medical School) cells were grown in Roswell Park Memorial Institute 1640 medium (RPMI) supplemented with 10%FBS 100 units/mL penicillin, 100 μg/mL streptomycin, and 25 ng/mL amphotericin B. A549 (CCL-185) and U-2 OS (HTB-96) were obtained from ATCC and grown in Dulbecco’s Modified Eagle Medium (DMEM) supplemented with the same concentrations of FBS and antibiotics as mentioned above. MCF10A (CRL-10317) were grown in DMEM/F-12 (Invitrogen #11330-032) media supplemented with 5% Horse serum (Invitrogen#16050-122), EGF (20ng/ml final), Hydrocortisone (0.5 mg/ml final), Cholera Toxin (100 ng/ml final), Insulin (10μg/ml final), 1% Pen/Strep (100x solution, Invitrogen #15070-063).

### Cell Treatments

For H_2_O_2_ treatments, H_2_O_2_ was diluted in PBS, then added directly to the media to get the final concentrations indicated in each experiment. The stock of H_2_O_2_ (Fisher Scientific AAL13235AP) was replaced monthly. Cells were treated with J14 (MedChemExpress, HY-135008) and added directly to the media for a final concentration of 20μM. A stock concentration of 35mM of Conoidin A (Cayman Chemical Item no. 15605) made in DMSO. The stock was diluted in media and then added to cells to obtain the indicated final concentration. Tert-butyl hydroperoxide (TBHP, 70% solution in water) was obtained from Acros Organics and diluted in PBS. The TBHP PBS stock was then added directly to media to obtain the indicated final concentration. A stock solution of Menadione (Sigma Aldrich M5625) was made in 100% Ethanol, diluted in media, and then added to cells to obtain the final concentration indicated for each experiment.

### Cell Line Construction

Lentivirus was produced and infection was carried out using protocol from Tiscornia et al., 2006. Transfection was carried out using the protocol from Mirus using the TransIT Transfection Reagent LT1. The SRXN1 overexpression vector was designed and constructed using VectorBuilder (https://en.vectorbuilder.com/). The vector is a lentiviral vector that expresses mCerulean-NLS-P2A-T2A-SRXN1 from the PGK promoter and harbors a blasticidin resistance gene. The NLS (nuclear localization sequence) tagged mCerulean is separated from SRXN1 with the P2A-T2A self-cleavage site so that SRXN1 is independent of the nuclear mCerulean signal. The mCerulean signal allows the verification that the construct is expressed in cells. MCF7 cells were infected with the lentivirus, cells were selected using blasticidin and clones were isolated and validated. The PRDX1 knockout line was made by using a CRISPR/Cas9 lentiviral vector from VectorBuilder (hPRDX1[gRNA#1077]). MCF7 cells were infected with lentivirus and selected using blasticidin. Individual clones were isolated and validated using Western Blot. SRXN1 knockdown cells were made by using plasmid from Vector Builder (pLV[shRNA]-Puro-U6>hSRXN1[shRNA#1]) to transfect MCF7 cells. Cells were then selected using puromycin and the knockdown was validated using Immunofluorescence.

### Immunofluorescence

Cells (1700 cells/well) were plated in glass bottom 96 multi-well plates (CellVis) or polystyrene plates from CellCarrier-96. Cells were allowed to attach for two days and treated at different time points. They were fixed with 2% PFA for 10 minutes, permeabilized using 0.1% Triton X-100 in PBS for 10mins, blocked with 2% BSA in PBS and incubated o/n at 4°C in primary antibodies made with 2% BSA and 0.1% Tween in PBS. The cells were then washed two times with PBS followed by incubation with secondary antibodies at room temperature for 1 hour. The cells were then washed two times in PBS followed by staining with DAPI and imaged in PBS. Images were analyzed Cell Profiler (McQuin et al., 2018). To obtain cytoplasmic levels of FOXO1, a ring of 3 pixels wide is was drawn around the nuclear mask and mean cytoplasmic FOXO1 was extracted using this mask. All plots were made using MATLAB. Primary antibodies used: Anti-FOXO1 (C29H4) from Cell Signaling Cat# 2880S (1:500), Anti-p53 (DO-1) from Santa Cruz Cat# sc-126 (1:500), Anti-Sulfiredoxin from Santa Cruz Cat# sc-166786 (1:100), Recombinant Anti-Peroxiredoxin 1/PAG antibody [EPR5433] (ab109498) (1:1000), NRF2 (D1Z9C) XP Rabbit mAb #12721 (1:500), NFAT1 (D43B1) XP Rabbit mAb #5861 (1:300), NF-kB p65 (D14E12) XP Rabbit mAb #8242 (1:400), Anti-HSF1 antibody 10H8 from Stress Marq Biosciences (1:200), c-Fos (9F6) Rabbit mAb # 2250 (1:1000), and c-Jun (60A8) Rabbit mAb #9165 (1:400) from Cell Signaling. Secondary Antibodies used: Alexa Fluor 488 goat anti-rabbit, Alexa Fluor 546 goat anti-rabbit, Alex Fluor 647 goat anti-mouse and Alexa Fluor 594 goat anti-mouse from Invitrogen. All secondary antibodies were used at a concentration of 1:500. DAPI was imaged using the AT-DAPI Filter Set (Chroma) for 40-60 ms. Alexa Fluor 488 was imaged using (Chroma) AT-EGFP/F Filter Set for 600-800 ms, Alexa Fluor 594 was imaged using (Chroma) AT-TR/mCH Filter Set for 600-800 ms, Alexa Fluor 546 was imaged using (Chroma) AT-TRITC/CY3 filter set and Alex Fluor 647 was imaged using AT-CY5.5 filter set.

### Live Cell Microscopy

Cells (15,000 cells / well) were plated on 12 well glass bottom plates (CellVis) which were coated with poly L-lysine (Sigma) and allowed to attach for 48hrs. The cells were grown in the appropriate media as mentioned above and then rinsed with PBS and given DMEM Fluorobrite (ThermoFisher) media with 2% FBS,100 units/mL penicillin, 100 μg/mL streptomycin, 25 ng/mL amphotericin B, and 1x Glutamax (ThermoFisher). Cells were imaged every 15-20 minutes for 24-48hrs by a Nikon Eclipse Ti-E microscope. Humidity, temperature at 37°C and 5% CO_2_ levels was maintained using the OKO labs incubation system. H2B-CFP was imaged using the C-FL AT ECFP/Cerulean Filter Set (Chroma) for 20–40 ms. FOXO1–mVenus was imaged using (Chroma) ET-EYFP Filter Set for 600-800 ms, p53-mCherry was imaged using (Chroma) AT-TR/mCH Filter Set for 600-800 ms. Movies were analyzed using single cell tracking software in MATLAB (Reyes et al., 2018)

### Western Blot

Cells (approx.100,000 cells/dish) were plated to a 6 cm dish and incubated for 48 hours. They were washed with PBS, scraped off the plate, centrifuged and the cell pellet was lysed using a Lysis Buffer (25 mM Tris pH 7.6, 150 mM NaCl, 1% NP-40, 1% Na-deoxycholate, and 0/1% SDS in water + protease inhibitor cocktail [Sigma] + phosphatase inhibitor cocktail [Sigma-Aldrich] + okadaic acid + sodium fluoride). The cells were spun down and the supernatant was used to measure protein concentration by using a Bradford Assay (BioRad). Equal protein concentrations were loaded onto a NuPAGE 4-12% Bis-Tris gels (Invitrogen). Protein was transferred to a nitrocellulose membrane and incubated in blocking solution (PBS, 5% BSA, 0.1% Tween 20) for 1 hour at room temperature. The membrane was incubated with primary antibodies at 4°C overnight, rinsed three times with PBST and incubated with secondary antibodies for 1 hour at room temperature. The blot was imaged on the LI-COR Odyssey. Primary Antibodies used: Anti-Peroxiredoxin-SO3 antibody (ab16830) (1:1000), Recombinant Anti-Peroxiredoxin 1/PAG antibody [EPR5433] (ab109498) (1:1000) from Abcam and actin (Sigma) was used for primary staining. Secondary antibodies used: 680LT secondary (LICOR-IRDye) and 800CW secondary (LICOR-IRDye), both at 1:10000 concentration.

### Single Cell ATAC and RNA sequencing

Chromium Next GEM Single Cell Multiome ATAC + Gene Expression kit (Product code: 1000285) from 10X Genomics was used for transposition, GEM generation and Barcoding, ATAC Library Construction, cDNA Amplication, Gene expression and library construction (CG000338). MCF7 cells were plated (50,000 cells/well) on plastic 6 well plates and allowed to attach for 48 hrs. Nuclei were isolated the cells treated with different concentrations of H_2_O_2_ for 5hrs using the Nuclei Isolation for Single Cell Multiome ATAC + Gene Expresssion protocol (CG000365) by standardizing the time of lysis to obtain a nuclei quality of A and B. All further steps with the kit were performed by the University of Arizona Genetics Core. Sequencing was carried out using Novogene.

### Bulk RNA sequencing

Cells (50,000 cells/well) were plated on plastic 6 well plates and allowed to attach for 2 days. Cells were then treated with different concentrations of H_2_O_2_ for 5hrs. RNA was isolated using RNeasy Mini Kit (Cat. No. 74104) from Qiagen. The isolated RNA was then sent to Novogene for sequencing.

### Analysis of Bulk RNA sequencing and single cell ATAC sequencing

#### Bulk Differential Expression Analysis

Analysis was performed by importing BAM files with the *Rsubread* package and *featureCounts* program dropping all genes with counts less than 10 across experimental conditions^67^. We then performed differential expression analysis on the dataset using *DEseq2* ^68^. J14, Sulfiredoxin, and wild-type samples were compared between 50μM hydrogen peroxide and no treatment. A list of genes was generated for each set of replicates from these comparisons using thresholds after effect size shrinkage using the *apeglm* package^69^. These lists were combined to evaluate overall expression changes across each treatment condition. We also evaluated expression differences that were of opposite direction across the J14 and sulfiredoxin conditions. We compared differential expression of untreated sulfiredoxin and J14 samples with their wild-type counterparts to get an estimate of expression differences from these conditions independent of hydrogen peroxide and confirmed these trends using Principal Component Analysis.

#### Bulk Gene Set Enrichment Analysis

Gene Set Enrichment Analysis was performed to evaluate differences in sulfiredoxin and J14 samples treated with 50μM hydrogen peroxide using the clusterProfiler R package^70–72^ (Yu et. al. 2012, Wu et. al. 2021) and functions from the enrichplot package [Yu G (2022) *enrichplot: Visualization of Functional Enrichment Result*. R package version 1.18.3, https://yulab-smu.top/biomedical-knowledge-mining-book/]. We included KEGG gene sets and custom gene sets of p53 and NRF2 target genes. The list of p53 target genes was obtained from Fisher, 2017^73^ and included p53 targets with direct regulation in seven different studies. NRF2 target genes were obtained from Okazaki et. al., 2020^74^.

#### Single Cell Sequencing Analysis

Fragment files generated via 10x Genomics were analyzed using the ArchR pipeline^75^. Untreated cells and those treated with 50uM and 75uM hydrogen peroxide were separated into six clusters using an iterative latent semantic indexing algorithm acting on tiles of 500-bp for each cell using the ArchR function *IterativeLSI* for dimensionality reduction and Seurat’s *findClusters* with a resolution of 0.14. Peaks were called in each cluster using MACS2 using the *addReproduciblePeakSet* function in ArchR. The *getMarkerFeatures* function was used to generate lists of relevant features from each of clusters 2, 3, and 6 with clusters with high numbers of untreated cells used as background groups (clusters 1, 4, and 5). Each cluster was analyzed to assess the prevalence of relevant transcription factors with JASPAR2020 binding profiles. Motif enrichment was evaluated on the generated features for each cluster for motifs with FDR < 0.01 and |LFC| > 1.3 using the *peakAnnoEnrichment* function in ArchR. ChromVAR deviations^76^ for each transcription factor were evaluated on a per-cell basis. Transcription factors were ordered by average ChromVAR deviations in each cluster and compared to z-scores of expression levels for each gene from scRNAseq data.

## Acknowledgments

We thank members of the Paek lab, the Thorne lab, J. Stewart-Ornstein and S. Chen for helpful comments and discussion. We also thank A. Capaldi and T. Weinert for feedback on the manuscript. This work was supported by National Institutes of Health Grants RO1GM130864 (A.P.) and NIH T32-GM132008 (W.M.S).

---

**Supplemental Figure 1:**
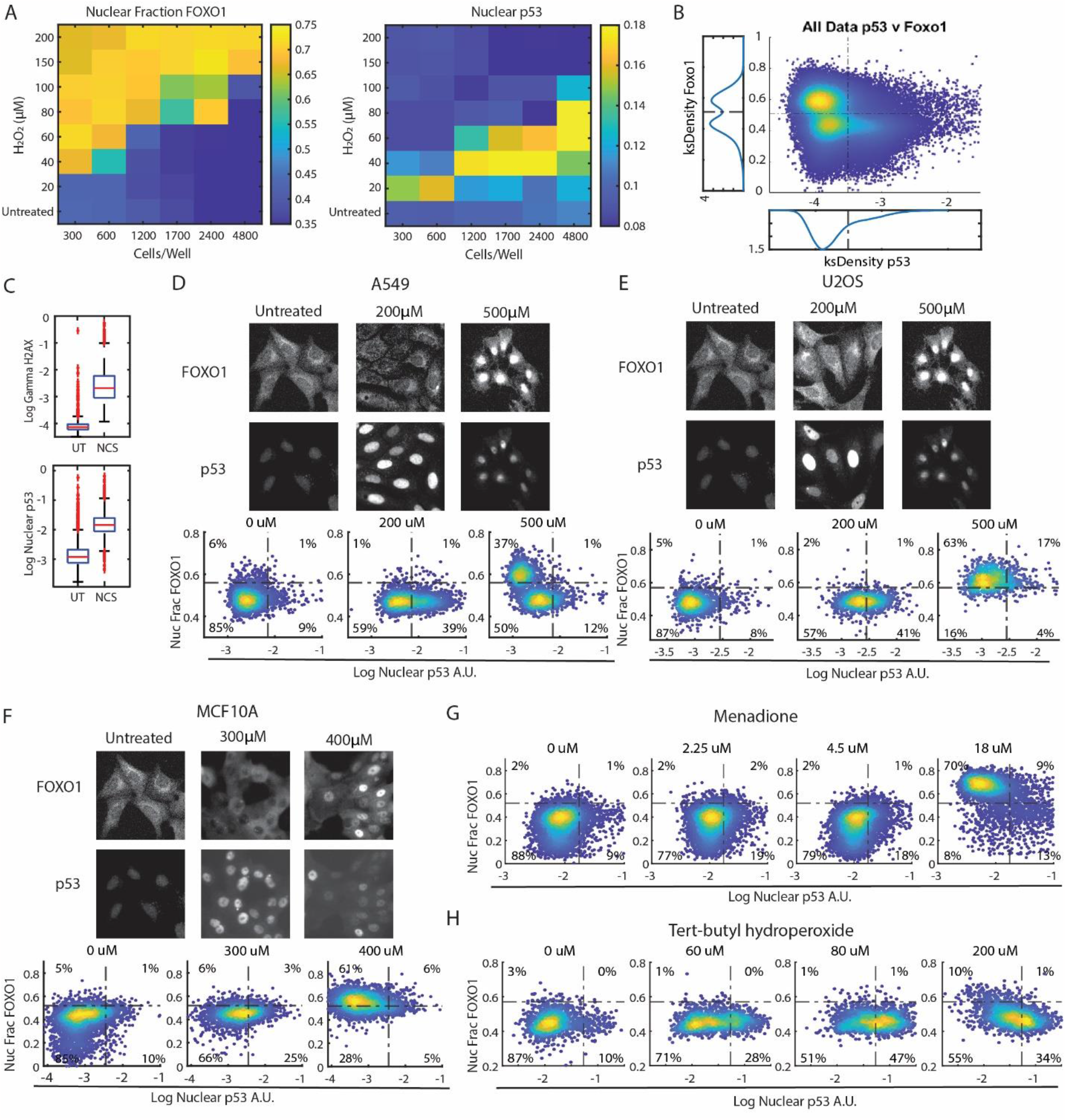
Other cell lines show mutually exclusive activity of FOXO1 and p53. (A) Heat maps showing the nuclear fraction of FOXO1 and nuclear p53 levels of cells plated at specified cell numbers (bottom) and indicated concentrations of H_2_O_2_ (left). (B) Density colored scatter plot of all cells used in Figure 2B to determine thresholds for the fraction of FOXO1 and nuclear levels of p53. Population density plots of nuclear fraction of FOXO1 (left side) and nuclear p53 (below) used for thresholding of the density plots shown in Figure 1B. (C) Bar and whisker plots of nuclear γH2AX and nuclear p53 levels after 3 hours of treatment with 400ng/ml of Neocarzinostatin (NCS). (D-F) Immunofluorescence images and density colored scatter plots of nuclear fraction of FOXO1 and nuclear p53 at the indicated concentrations of H_2_O_2_ for 5 hours for (D) A549 (E) U2OS and (F) MCF10A cell lines. (G and H) Density colored scatter of cells treated with indicated concentrations of Menadione and tert-butyl hydroperoxide (TBHP) for 5 hours.

**Supplemental Fig 2:**
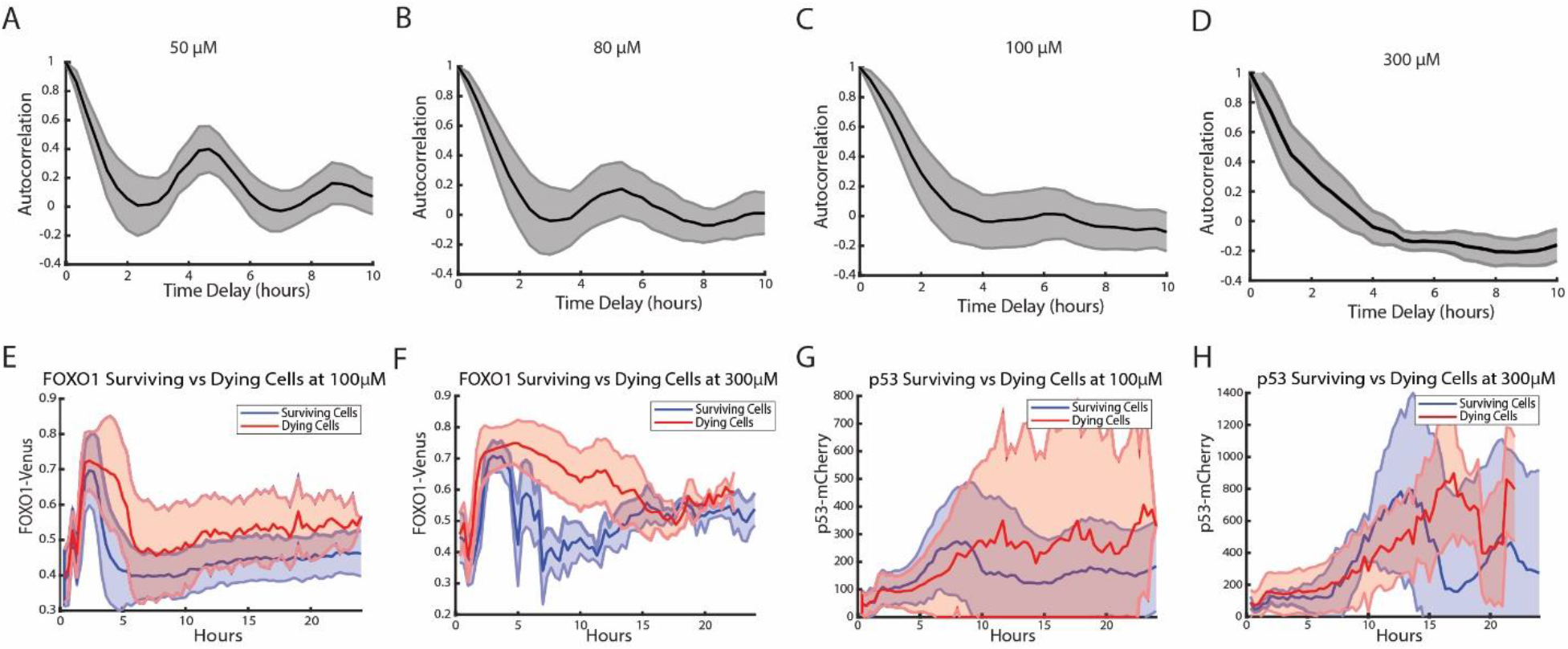
p53 dynamics in response to H_2_O_2_ stress. (A) Autocorrelation of p53 trajectories of cells treated with 50μM of H_2_O_2_ from Figure 2B. (B) Autocorrelation of p53 trajectories after FOXO1 phase of cells treated with 80μM of H_2_O_2_ from Figure 2C (C) Autocorrelation of p53 trajectories after FOXO1 phase of cells treated with 100μM of H_2_O_2_ from Figure 2D. (D) Autocorrelation of p53 trajectories after FOXO1 phase of cells treated with 300μM of H_2_O_2_ from Figure 2E. (E) FOXO1-mVenus levels in surviving (blue) and dead (red) cells treated with 100μM of H_2_O_2_ over 24 hours. (F) FOXO1-mVenus levels in surviving (blue) and dead (red) cells treated with 300μM of H_2_O_2_ over 24 hours. (G) p53-mCherry levels in surviving (blue) and dead (red) cells treated with 100μM of H_2_O_2_ over 24 hours. (H) p53-mCherry levels in surviving (blue) and dead (red) cells treated with 100μM of H_2_O_2_ over 24 hours.

**Supplemental Fig 3:**
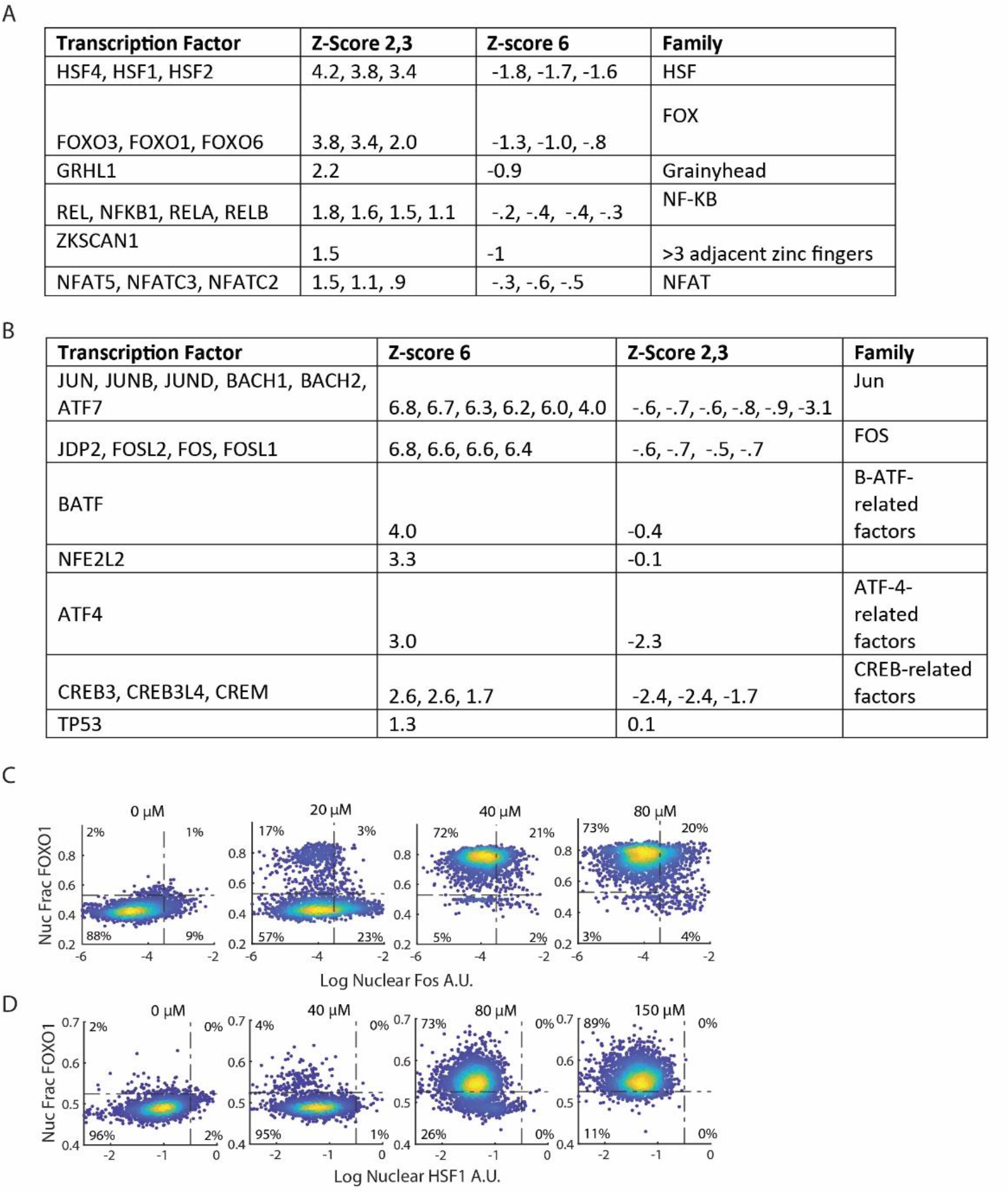
Other Transcription factors are activated with FOXO1 and p53 in response to oxidative stress. (A) TF binding motifs enriched in clusters 2 and 3 of single-cell ATAC data. Motifs of TFs not expressed in gene expression data (>.1 counts per cell) were omitted. (B) TF binding motifs enriched in cluster 6 of single-cell ATAC data. Motifs of TFs not expressed in gene expression data (>.1 counts per cell) were omitted. (C) Density colored scatter plots of nuclear fraction of FOXO1 and log of nuclear levels of FOA for cells treated at indicated concentrations of H_2_O_2_ for 5 hours. (D) Density colored scatter plots of nuclear fraction of FOXO1 and nuclear fraction of HSF1 for cells treated with indicated concentrations of H_2_O_2_ for 5 hours.

**Supplemental Figure 4:**
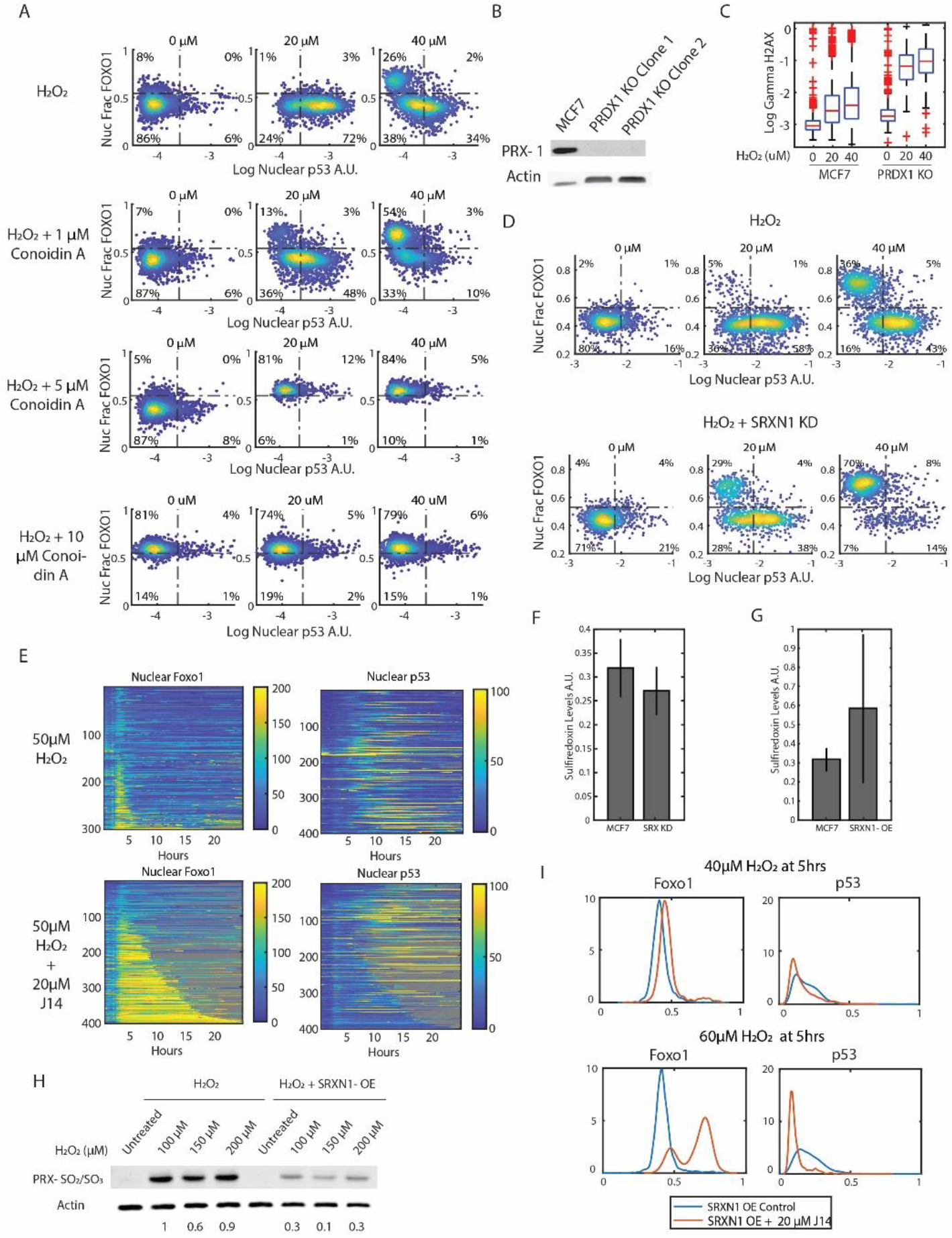
Overexpression/Inhibition of Sulfiredoxin controls the activation of FOXO1 and p53. (A) Density colored scatter plots representing the nuclear fraction of FOXO1 and nuclear p53 levels of cells treated with H_2_O_2_ alone (top) or with 1 μM (2^nd^ from top), 5μM (3^rd^ from top) and 10 μM (4^th^ from top) of Conoidin A at indicated concentrations of H_2_O_2_ for 5 hours. (B) Western Blot stained with PRDX1 antibody to validate PRDX1 knockout in MCF7 cells. (C) Bar and whisker plots of gamma H2AX levels in MCF7 cells and PRDX1 knockout cells treated with 20μM of H_2_O_2_ for 3 hours. (D) Density scatter plots of cells treated with H_2_O_2_ as a control for cells with a SRXN1 knockdown treated with indicated concentrations of H_2_O_2_ for 3 hours. (E) Heatmaps of single cell traces of the nuclear fraction of FOXO1-mVenus (left) and nuclear p53-mCherry (right) in cells treated with 50μM H_2_O_2_ (top, n=306, 13% cell death) and 50μM H_2_O_2_ + 20μM J14 (n=406, 70% cell death) for 24 hours. Each row of the heat maps is a single cell over time. Both FOXO1 and p53 heat maps were sorted by the duration that FOXO1-mVenus remained in the nucleus. Gray indicates cell death. (F) Bar graphs of mean Sulfiredoxin levels in cells with the shRNA knockdown compared to the MCF7 control. (G) Bar graphs of mean levels of Sulfiredoxin in cells with SRXN1 overexpression compared to MCF7 cells using immunofluorescence. (H) Western blot of MCF7 cells and Sulfiredoxin (SRXN1) overexpressing cells treated with indicated concentrations of H_2_O_2_ for 3 hours and stained for hyperoxidized (SO_2_/SO_3_) PRDX1/2 and Actin. (I) Population density plots of nuclear fraction of FOXO1 and nuclear p53 levels in cells overexpressing Sulfiredoxin (SRXN1-OE) after treatment with H_2_O_2_ and the same cells treated with 20μM of J14 at indicated concentrations of H_2_O_2_. Blue line indicates cells with overexpression of Sulfiredoxin and the red line indicates cells overexpressing Sulfiredoxin treated with 20μM J14.

**Supplemental Figure 5.**
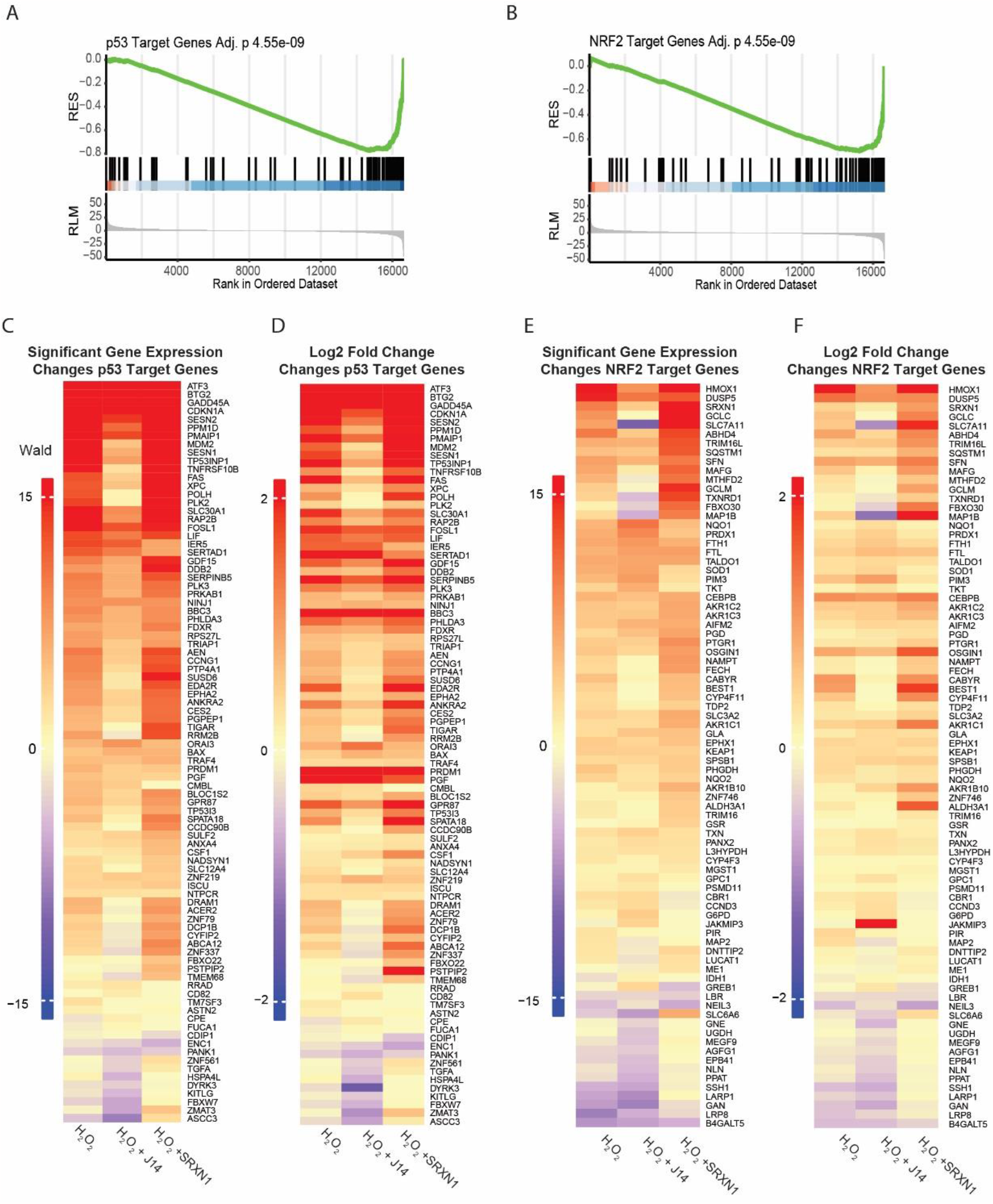
Differential gene expression analysis of p53 and NRF2 target genes. (A) GSEA of p53 target genes in J14 + H_2_O_2_ samples as compared to SRXN1-OE + H_2_O_2_ samples (B) GSEA of NRF2 target genes in J14 + H_2_O_2_ samples as compared to SRXN1-OE + H_2_O_2_ samples. (C) Wald statistic and (D) log_2_ fold changes of p53 target genes in H_2_O_2_ treated cells vs. PBS controls, H_2_O_2_ + J14 treated cells vs J14 treated controls and H_2_O_2_ treated SRXN1-OE cells vs. SRXN1-OE PBS controls. (E) Wald statistic and (F) log_2_ fold changes of NRF2 target genes in H_2_O_2_ treated cells vs. PBS controls, H_2_O_2_ + J14 treated cells vs J14 treated controls and H_2_O_2_ treated SRXN1-OE cells vs. SRXN1-OE PBS controls. H_2_O_2_ concentration is 50μM, J14 20μM.

**Supplemental Movie 1**: Example movie showing FOXO1-mVenus and p53-mCherry activity after treatment with 50μM of H_2_O_2_ in MCF7 cells for a period of 24hrs.H_2_O_2_ was added at the 20-minute timepoint.

**Supplemental Movie 2**: Example movie showing FOXO1-mVenus and p53-mCherry activity after treatment with 80μM of H_2_O_2_ in MCF7 cells for a period of 24hrs.H_2_O_2_ was added at the 20-minute timepoint.

**Supplemental Movie 3**: Example movie showing FOXO1-mVenus and p53-mCherry activity after treatment with 100μM of H_2_O_2_ in MCF7 cells for a period of 24hrs.H_2_O_2_ was added at the 20-minute timepoint.

**Supplemental Movie 4**: Example movie showing FOXO1-mVenus and p53-mCherry activity after treatment with 300μM of H_2_O_2_ in MCF7 cells for a period of 24hrs.H_2_O_2_ was added at the 20-minute timepoint.

## References

1. Sies, H., and Jones, D.P. (2020). Reactive oxygen species (ROS) as pleiotropic physiological signalling agents. Nat. Rev. Mol. Cell Biol. 2020 217 21, 363–383. 10.1038/s41580-020-0230-3.

2. Arnold, R.S., Shi, J., Murad, E., Whalen, A.M., Sun, C.Q., Polavarapu, R., Parthasarathy, S., Petros, J.A., and Lambeth, J.D. (2001). Hydrogen peroxide mediates the cell growth and transformation caused by the mitogenic oxidase Nox1. Proc. Natl. Acad. Sci. U. S. A. 98, 5550–5555. 10.1073/PNAS.101505898/ASSET/841657EE-35FA-4418-B072-34FF4F5FB357/ASSETS/GRAPHIC/PQ1015058004.JPEG.

3. Sigaud, S., Evelson, P., and González-Flecha, B. (2004). H2O2-Induced Proliferation of Primary Alveolar Epithelial Cells Is Mediated by MAP Kinases. https://home.liebertpub.com/ars 7, 6–13. 10.1089/ARS.2005.7.6.

4. Niethammer, P., Grabher, C., Look, A.T., and Mitchison, T.J. (2009). A tissue-scale gradient of hydrogen peroxide mediates rapid wound detection in zebrafish. Nature 459, 996. 10.1038/NATURE08119.

5. Manford, A.G., Rodríguez-Pérez, F., Shih, K.Y., Shi, Z., Berdan, C.A., Choe, M., Titov, D. V., Nomura, D.K., and Rape, M. (2020). A Cellular Mechanism to Detect and Alleviate Reductive Stress. Cell 183, 46-61.e21. 10.1016/J.CELL.2020.08.034/ATTACHMENT/F5F9EB00-D5C4-470D-B2E2-82BA3A590A99/MMC1.PDF.

6. Sena, L.A., and Chandel, N.S. (2012). Physiological roles of mitochondrial reactive oxygen species. Mol. Cell 48, 158–167. 10.1016/J.MOLCEL.2012.09.025.

7. Thomas, C., Mackey, M.M., Diaz, A.A., and Cox, D.P. (2009). Hydroxyl radical is produced via the Fenton reaction in submitochondrial particles under oxidative stress: Implications for diseases associated with iron accumulation. Redox Rep. 14, 102–108. 10.1179/135100009X392566.

8. Keyer, K., and Imlay, J.A. (1996). Superoxide accelerates DNA damage by elevating free-iron levels. Proc. Natl. Acad. Sci. U. S. A. 93, 13635–13640. 10.1073/pnas.93.24.13635.

9. Pravda, J. Hydrogen peroxide and disease: towards a unified system of pathogenesis and therapeutics. 10.1186/s10020-020-00165-3.

10. Marinho, H.S., Real, C., Cyrne, L., Soares, H., and Antunes, F. (2014). Hydrogen peroxide sensing, signaling and regulation of transcription factors. Redox Biol. 2, 535–562. 10.1016/j.redox.2014.02.006.

11. Liu, D., and Xu, Y. (2011). P53, oxidative stress, and aging. Antioxidants Redox Signal. 15, 1669–1678. 10.1089/ars.2010.3644.

12. Madan, E., Gogna, R., Bhatt, M., Pati, U., Kuppusamy, P., and Mahdi, A.A. (2011). Regulation of glucose metabolism by p53: Emerging new roles for the tumor suppressor. Oncotarget 2, 948–957. 10.18632/oncotarget.389.

13. Klotz, L.O., Sánchez-Ramos, C., Prieto-Arroyo, I., Urbánek, P., Steinbrenner, H., and Monsalve, M. (2015). Redox regulation of FoxO transcription factors. Redox Biol. 6, 51. 10.1016/J.REDOX.2015.06.019.

14. Shi, T., and Dansen, T.B. (2020). Reactive Oxygen Species Induced p53 Activation: DNA Damage, Redox Signaling, or Both? Antioxidants Redox Signal. 33, 839–859. 10.1089/ars.2020.8074.

15. Ma, Q. (2013). Role of Nrf2 in oxidative stress and toxicity. Annu. Rev. Pharmacol. Toxicol. 53, 401– 426. 10.1146/annurev-pharmtox-011112-140320.

16. Ahn, S.G., and Thiele, D.J. (2003). Redox regulation of mammalian heat shock factor 1 is essential for Hsp gene activation and protection from stress. Genes Dev. 17, 516–528. 10.1101/gad.1044503.

17. Lin, T.-Y., Cantley, L.C., and DeNicola, G.M. (2016). NRF2 Rewires Cellular Metabolism to Support the Antioxidant Response. In A Master Regulator of Oxidative Stress - The Transcription Factor Nrf2 (InTech). 10.5772/65141.

18. Shi, T., van Soest, D.M.K., Polderman, P.E., Burgering, B.M.T., and Dansen, T.B. (2021). DNA damage and oxidant stress activate p53 through differential upstream signaling pathways. Free Radic. Biol. Med. 172, 298–311. 10.1016/J.FREERADBIOMED.2021.06.013.

19. Lehtinen, M.K., Yuan, Z., Boag, P.R., Yang, Y., Villén, J., Becker, E.B.E., DiBacco, S., de la Iglesia, N., Gygi, S., Blackwell, T.K., et al. (2006). A Conserved MST-FOXO Signaling Pathway Mediates Oxidative-Stress Responses and Extends Life Span. Cell 125, 987–1001. 10.1016/j.cell.2006.03.046.

20. Veal, E.A., Day, A.M., and Morgan, B.A. (2007). Hydrogen Peroxide Sensing and Signaling. Mol. Cell 26, 1–14. 10.1016/J.MOLCEL.2007.03.016.

21. Sies, H. (2018). On the history of oxidative stress: Concept and some aspects of current development. Curr. Opin. Toxicol. 7, 122–126. 10.1016/J.COTOX.2018.01.002.

22. Quinn, J., Findlay, V.J., Dawson, K., Millar, J.B.A., Jones, N., Morgan, B.A., and Toone, W.M. (2002). Distinct Regulatory Proteins Control the Graded Transcriptional Response to Increasing H2O2 Levels in Fission Yeast Schizosaccharomyces pombe. Mol. Biol. Cell 13, 805. 10.1091/MBC.01-06-0288.

23. Eshaghi, M., Lee, J.H., Zhu, L., Poon, S.Y., Li, J., Cho, K.H., Chu, Z., Karuturi, R.K.M., and Liu, J. (2010). Genomic Binding Profiling of the Fission Yeast Stress-Activated MAPK Sty1 and the bZIP Transcriptional Activator Atf1 in Response to H2O2. PLoS One 5. 10.1371/JOURNAL.PONE.0011620.

24. Vivancos, A.P., Castillo, E.A., Jones, N., Ayté, J., and Hidalgo, E. (2004). Activation of the redox sensor Pap1 by hydrogen peroxide requires modulation of the intracellular oxidant concentration. Mol. Microbiol. 52, 1427–1435. 10.1111/J.1365-2958.2004.04065.X.

25. Bozonet, S.M., Findlay, V.J., Day, A.M., Cameron, J., Veal, E.A., and Morgan, B.A. (2005). Oxidation of a eukaryotic 2-Cys peroxiredoxin is a molecular switch controlling the transcriptional response to increasing levels of hydrogen peroxide. J. Biol. Chem. 280, 23319–23327. 10.1074/jbc.M502757200.

26. Calvo, I.A., Boronat, S., Domènech, A., García-Santamarina, S., Ayté, J., and Hidalgo, E. (2013). Dissection of a redox relay: H2O2-dependent activation of the transcription factor Pap1 through the peroxidatic Tpx1-thioredoxin cycle. Cell Rep. 5, 1413–1424. 10.1016/J.CELREP.2013.11.027.

27. Day, A.M., Brown, J.D., Taylor, S.R., Rand, J.D., Morgan, B.A., and Veal, E.A. (2012). Inactivation of a Peroxiredoxin by Hydrogen Peroxide Is Critical for Thioredoxin-Mediated Repair of Oxidized Proteins and Cell Survival. Mol. Cell 45, 398–408. 10.1016/j.molcel.2011.11.027.

28. Brown, J.D., Day, A.M., Taylor, S.R., Tomalin, L.E., Morgan, B.A., and Veal, E.A. (2013). A peroxiredoxin promotes H2O2 signaling and oxidative stress resistance by oxidizing a thioredoxin family protein. Cell Rep. 5, 1425–1435. 10.1016/J.CELREP.2013.10.036.

29. Castillo, E.A., Ayté, J., Chiva, C., Moldón, A., Carrascal, M., Abián, J., Jones, N., and Hidalgo, E. (2002). Diethylmaleate activates the transcription factor Pap1 by covalent modification of critical cysteine residues. Mol. Microbiol. 45, 243–254. 10.1046/J.1365-2958.2002.03020.X.

30. Vivancos, A.P., Castillo, E.A., Biteau, B., Nicot, C., Ayté, J., Toledano, M.B., and Hidalgo, E. (2005). A cysteine-sulfinic acid in peroxiredoxin regulates H2O 2-sensing by the antioxidant Pap1 pathway. Proc. Natl. Acad. Sci. U. S. A. 102, 8875–8880. 10.1073/pnas.0503251102.

31. Stöcker, S., Maurer, M., Ruppert, T., and Dick, T.P. (2018). A role for 2-Cys peroxiredoxins in facilitating cytosolic protein thiol oxidation. Nat. Chem. Biol. 14, 148–155. 10.1038/nchembio.2536.

32. Sobotta, M.C., Liou, W., Stöcker, S., Talwar, D., Oehler, M., Ruppert, T., Scharf, A.N.D., and Dick, T.P. (2015). Peroxiredoxin-2 and STAT3 form a redox relay for H2O2 signaling. Nat. Chem. Biol. 11, 64–70. 10.1038/nchembio.1695.

33. Talwar, D., Messens, J., and Dick, T.P. (2020). A role for annexin A2 in scaffolding the peroxiredoxin 2– STAT3 redox relay complex. Nat. Commun. 2020 111 11, 1–11. 10.1038/s41467-020-18324-9.

34. Putker, M., Vos, H.R., Van Dorenmalen, K., De Ruiter, H., Duran, A.G., Snel, B., Burgering, B.M.T., Vermeulen, M., and Dansen, T.B. (2015). Evolutionary Acquisition of Cysteines Determines FOXO Paralog-Specific Redox Signaling. Antioxid. Redox Signal. 22, 15. 10.1089/ARS.2014.6056.

35. Hopkins, B.L., Nadler, M., Skoko, J.J., Bertomeu, T., Pelosi, A., Shafaei, P.M., Levine, K., Schempf, A., Pennarun, B., Yang, B., et al. (2018). A Peroxidase Peroxiredoxin 1-Specific Redox Regulation of the Novel FOXO3 microRNA Target let-7. Antioxid. Redox Signal. 28, 62. 10.1089/ARS.2016.6871.

36. Brunet, A., Bonni, A., Zigmond, M.J., Lin, M.Z., Juo, P., Hu, L.S., Anderson, M.J., Arden, K.C., Blenis, J., and Greenberg, M.E. (1999). Akt Promotes Cell Survival by Phosphorylating and Inhibiting a Forkhead Transcription Factor.

37. Brooks, C.L., and Gu, W. (2011). p53 Regulation by Ubiquitin. FEBS Lett. 585, 2803. 10.1016/J.FEBSLET.2011.05.022.

38. Cooke, M.S., Evans, M.D., Dizdaroglu, M., and Lunec, J. (2003). Oxidative DNA damage: mechanisms, mutation, and disease. FASEB J. 17, 1195–1214. 10.1096/FJ.02-0752REV.

39. Lasick, K.A., Jose, E., Samayoa, A.M., Shanks, L., Pond, K.W., Thorne, C.A., and Paek, A.L. (2023). FOXO nuclear shuttling dynamics are stimulus-dependent and correspond with cell fate. Mol. Biol. Cell 34, ar21. 10.1091/MBC.E22-05-0193/ASSET/IMAGES/LARGE/MBC-34-AR21-G005.JPEG.

40. Hanson, R.L., and Batchelor, E. (2022). Coordination of MAPK and p53 dynamics in the cellular responses to DNA damage and oxidative stress. Mol. Syst. Biol. 18, 11401. 10.15252/MSB.202211401.

41. Lahav, G., Rosenfeld, N., Sigal, A., Geva-Zatorsky, N., Levine, A.J., Elowitz, M.B., and Alon, U. (2004). Dynamics of the p53-Mdm2 feedback loop in individual cells. Nat. Genet. 10.1038/ng1293.

42. Batchelor, E., Loewer, A., Mock, C., and Lahav, G. (2011). Stimulus-dependent dynamics of p53 in single cells. Mol. Syst. Biol. 7, 1–8. 10.1038/msb.2011.20.

43. Gaglia, G., Rashid, R., Yapp, C., Joshi, G.N., Li, C.G., Lindquist, S.L., Sarosiek, K.A., Whitesell, L., Sorger, P.K., and Santagata, S. (2020). HSF1 phase transition mediates stress adaptation and cell fate decisions. Nat. Cell Biol. 22, 151. 10.1038/S41556-019-0458-3.

44. Fornes, O., Castro-Mondragon, J.A., Khan, A., Van Der Lee, R., Zhang, X., Richmond, P.A., Modi, B.P., Correard, S., Gheorghe, M., Baranašić, D., et al. (2020). JASPAR 2020: update of the open-access database of transcription factor binding profiles. Nucleic Acids Res. 48, D87. 10.1093/NAR/GKZ1001.

45. Perkins, A., Nelson, K.J., Parsonage, D., Poole, L.B., and Karplus, P.A. Peroxiredoxins: Guardians Against Oxidative Stress and Modulators of Peroxide Signaling. 10.1016/j.tibs.2015.05.001.

46. Stöcker, S., Van Laer, K., Mijuskovic, A., and Dick, T.P. (2018). The Conundrum of Hydrogen Peroxide Signaling and the Emerging Role of Peroxiredoxins as Redox Relay Hubs. Antioxidants Redox Signal. 28, 558–573. 10.1089/ARS.2017.7162/ASSET/IMAGES/LARGE/FIGURE6.JPEG.

47. Haraldsen, J.D., Liu, G., Botting, C.H., Walton, J.G.A., Storm, J., Phalen, T.J., Kwok, L.Y., Soldati-Favre, D., Heintz, N.H., Müller, S., et al. (2009). IDENTIFICATION OF CONOIDIN A AS A COVALENT INHIBITOR OF PEROXIREDOXIN II. Org. Biomol. Chem. 7, 3040. 10.1039/B901735F.

48. Nguyen, J.B., Pool, C.D., Wong, C.Y.B., Treger, R.S., Williams, D.L., Cappello, M., Lea, W.A., Simeonov, A., Vermeire, J.J., and Modis, Y. (2013). Peroxiredoxin-1 from the human hookworm Ancylostoma ceylanicum forms a stable oxidized decamer and is covalently inhibited by conoidin A. Chem. Biol. 20, 991. 10.1016/J.CHEMBIOL.2013.06.011.

49. Skoko, J.J., Cao, J., Gaboriau, D., Attar, M., Asan, A., Hong, L., Paulsen, C.E., Ma, H., Liu, Y., Wu, H., et al. (2022). Redox regulation of RAD51 Cys319 and homologous recombination by peroxiredoxin 1. Redox Biol. 56, 102443. 10.1016/J.REDOX.2022.102443.

50. Jang, H.H., Lee, K.O., Chi, Y.H., Jung, B.G., Park, S.K., Park, J.H., Lee, J.R., Lee, S.S., Moon, J.C., Yun, J.W., et al. (2004). Two enzymes in one: Two yeast peroxiredoxins display oxidative stress-dependent switching from a peroxidase to a molecular chaperone function. Cell 117, 625–635. 10.1016/j.cell.2004.05.002.

51. Molin, M., Yang, J., Hanzén, S., Toledano, M.B., Labarre, J., and Nyström, T. (2011). Life span extension and H(2)O(2) resistance elicited by caloric restriction require the peroxiredoxin Tsa1 in Saccharomyces cerevisiae. Mol. Cell 43, 823–833. 10.1016/J.MOLCEL.2011.07.027.

52. Troussicot, L., Burmann, B.M., and Molin, M. (2021). Structural determinants of multimerization and dissociation in 2-Cys peroxiredoxin chaperone function. Structure 29, 640–654. 10.1016/J.STR.2021.04.007.

53. Shcherbik, N., and Pestov, D.G. (2019). The Impact of Oxidative Stress on Ribosomes: From Injury to Regulation. Cells 8. 10.3390/CELLS8111379.

54. Murphy, M.P. (2009). How mitochondria produce reactive oxygen species. Biochem. J. 417, 1–13. 10.1042/BJ20081386.

55. Bolduc, J.A., Nelson, K.J., Haynes, A.C., Lee, J., Reisz, J.A., Graff, A.H., Clodfelter, J.E., Parsonage, D., Poole, L.B., Furdui, C.M., et al. (2018). Novel hyperoxidation resistance motifs in 2-Cys peroxiredoxins. J. Biol. Chem. 293, 11901–11912. 10.1074/JBC.RA117.001690.

56. Portillo-Ledesma, S., Randall, L.M., Parsonage, D., Rizza, J.D., Andrew Karplus, P., Poole, L.B., Denicola, A., and Ferrer-Sueta, G. (2018). Differential kinetics of two-cysteine peroxiredoxin disulfide formation reveal a novel model for peroxide sensing. Biochemistry 57, 3416–3424. 10.1021/ACS.BIOCHEM.8B00188/SUPPL_FILE/BI8B00188_SI_001.PDF.

57. Randall, L.M., Ferrer-Sueta, G., and Denicola, A. (2013). Peroxiredoxins as Preferential Targets in H2O2-Induced Signaling. Methods Enzymol. 527, 41–63. 10.1016/B978-0-12-405882-8.00003-9.

58. Akter, S., Fu, L., Jung, Y., Lo Conte, M., Reed Lawson, J., Todd Lowther, W., Sun, R., Liu, K., Yang, J., and Carroll, K.S. (2018). Chemical Proteomics Reveals New Targets of Cysteine Sulfinic Acid Reductase. Nat. Chem. Biol. 14, 995. 10.1038/S41589-018-0116-2.

59. Delaunay, A., Pflieger, D., Barrault, M.B., Vinh, J., and Toledano, M.B. (2002). A Thiol Peroxidase Is an H2O2 Receptor and Redox-Transducer in Gene Activation. Cell 111, 471–481. 10.1016/S0092-8674(02)01048-6.

60. Bellezza, I., Giambanco, I., Minelli, A., and Donato, R. (2018). Nrf2-Keap1 signaling in oxidative and reductive stress. Biochim. Biophys. Acta - Mol. Cell Res. 1865, 721–733. 10.1016/J.BBAMCR.2018.02.010.

61. Fourquet, S., Guerois, R., Biard, D., and Toledano, M.B. (2010). Activation of NRF2 by Nitrosative Agents and H2O2 Involves KEAP1 Disulfide Formation. J. Biol. Chem. 285, 8463. 10.1074/JBC.M109.051714.

62. Nadeau, P.J., Charette, S.J., Toledano, M.B., and Landry, J. (2007). Disulfide Bond-mediated Multimerization of Ask1 and Its Reduction by Thioredoxin-1 Regulate H 2 O 2-induced c-Jun NH 2-terminal Kinase Activation and Apoptosis. Mol. Biol. Cell 18, 3903–3913. 10.1091/mbc.E07-05-0491.

63. Nadeau, P.J., Charette, S.J., and Landry, J. (2009). REDOX Reaction at ASK1-Cys250 Is Essential for Activation of JNK and Induction of Apoptosis. Mol. Biol. Cell 20, 3628. 10.1091/MBC.E09-03-0211.

64. Sun, X.Z., Vinci, C., Makmura, L., Han, S., Tran, D., Nguyen, J., Hamann, M., Grazziani, S., Sheppard, S., Gutova, M., et al. (2004). Formation of Disulfide Bond in p53 Correlates with Inhibition of DNA Binding and Tetramerization. https://home.liebertpub.com/ars 5, 655–665. 10.1089/152308603770310338.

65. Martín, D., Salinas, M., Fujita, N., Tsuruo, T., and Cuadrado, A. (2002). Ceramide and reactive oxygen species generated by H2O2 induce caspase-3-independent degradation of Akt/protein kinase B. J. Biol. Chem. 277, 42943–42952. 10.1074/jbc.M201070200.

66. Murata, H., Ihara, Y., Nakamura, H., Yodoi, J., Sumikawa, K., and Kondo, T. (2003). Glutaredoxin Exerts an Antiapoptotic Effect by Regulating the Redox State of Akt. J. Biol. Chem. 278, 50226–50233. 10.1074/jbc.M310171200.

67. Liao, Y., Smyth, G.K., and Shi, W. (2014). featureCounts: an efficient general purpose program for assigning sequence reads to genomic features. Bioinformatics 30, 923–930. 10.1093/BIOINFORMATICS/BTT656.

68. Love, M.I., Huber, W., and Anders, S. (2014). Moderated estimation of fold change and dispersion for RNA-seq data with DESeq2. Genome Biol. 15, 1–21. 10.1186/S13059-014-0550-8/FIGURES/9.

69. Zhu, A., Ibrahim, J.G., and Love, M.I. (2019). Heavy-tailed prior distributions for sequence count data: removing the noise and preserving large differences. Bioinformatics 35, 2084–2092. 10.1093/BIOINFORMATICS/BTY895.

70. Yu, G., Wang, L.G., Han, Y., and He, Q.Y. (2012). ClusterProfiler: An R package for comparing biological themes among gene clusters. Omi. A J. Integr. Biol. 16, 284–287. 10.1089/omi.2011.0118.

71. Wu, T., Hu, E., Xu, S., Chen, M., Guo, P., Dai, Z., Feng, T., Zhou, L., Tang, W., Zhan, L., et al. (2021). clusterProfiler 4.0: A universal enrichment tool for interpreting omics data. Innov. (Cambridge 2. 10.1016/J.XINN.2021.100141.

72. Subramanian, A., Tamayo, P., Mootha, V.K., Mukherjee, S., Ebert, B.L., Gillette, M.A., Paulovich, A., Pomeroy, S.L., Golub, T.R., Lander, E.S., et al. (2005). Gene set enrichment analysis: A knowledge-based approach for interpreting genome-wide expression profiles. Proc. Natl. Acad. Sci. U. S. A. 102, 15545–15550. 10.1073/PNAS.0506580102/SUPPL_FILE/06580FIG7.JPG.

73. Fischer, M. (2017). Census and evaluation of p53 target genes. Oncogene 2017 3628 36, 3943–3956. 10.1038/onc.2016.502.

74. Okazaki, K., Anzawa, H., Liu, Z., Ota, N., Kitamura, H., Onodera, Y., Alam, M.M., Matsumaru, D., Suzuki, T., Katsuoka, F., et al. (2020). Enhancer remodeling promotes tumor-initiating activity in NRF2-activated non-small cell lung cancers. Nat. Commun. 2020 111 11, 1–19. 10.1038/s41467-020-19593-0.

75. Granja, J.M., Corces, M.R., Pierce, S.E., Bagdatli, S.T., Choudhry, H., Chang, H.Y., and Greenleaf, W.J. (2021). Author Correction: ArchR is a scalable software package for integrative single-cell chromatin accessibility analysis (Nature Genetics, (2021), 53, 3, (403-411), 10.1038/s41588-021-00790-6). Nat. Genet. 53, 935. 10.1038/s41588-021-00850-x.

76. Schep, A.N., Wu, B., Buenrostro, J.D., and Greenleaf, W.J. (2017). chromVAR: Inferring transcription factor-associated accessibility from single-cell epigenomic data. Nat. Methods 14, 975. 10.1038/NMETH.4401.

